# Layered Single-Cell Heterogeneity in Hormone Receptor Signaling Across Mouse Organoids and Human ERα+ Cancer Cells

**DOI:** 10.64898/2026.03.19.712456

**Authors:** Pelin Yasar, Christopher R. Day, Brian D. Bennett, Jackson A. Hoffman, Laura G. Kammel, Trevor K. Archer, Joseph Rodriguez

## Abstract

Hormone receptor signaling is often interpreted through receptor abundance as a proxy for hormone responsiveness, yet the determinants of single-cell variability in hormone response remain unclear. Using murine mammary organoids, we mapped estrogen (E2) and progesterone (P4) responses at single-cell resolution. Despite controlled 3D culture conditions, basal cell-derived organoids exhibit striking variability in hormone-induced transcriptional responses, with ERα⁺ cells varying in the fraction of responsive genes engaged. This variability is not explained by receptor abundance alone. Instead, response magnitude correlates with expression of transcriptional co-regulators including *Ncoa1*, *Ncor1*, and *Ncor2*, suggesting that co-regulator balance contributes to variation in endocrine response magnitude. Organoids exhibit a mixed basal-luminal enhancer landscape and growth factor-dependent remodeling of ERα and PR protein abundance. MCF7 cells show delayed activation kinetics and reach a lower response plateau. Together, these findings reveal that hormone response magnitude varies independently of receptor abundance in mammary organoids and follows distinct activation dynamics in human ERα^+^ cancer cells.

## Introduction

Mammary gland development and remodeling depend on ovarian steroid hormones, primarily estrogen and progesterone. These hormones act through their cognate nuclear receptors estrogen receptor alpha (ERα) and progesterone receptor (PR)^1–3^. These receptors orchestrate gene expression programs that govern epithelial proliferation, differentiation, and morphogenesis within a bilayered epithelium composed of basal myoepithelial cells, luminal hormone-sensing and secretory cells, and an instructive stromal compartment^1,2,4^. Although ERα⁺ cells comprise only a subset of luminal epithelial cells, they function as hormone sensors that translate systemic endocrine signals into local paracrine cues controlling neighboring epithelial and stromal populations^5–7^. This sensor role, together with the tight spatial and temporal regulation of ERα and PR expression, suggests that hormone signaling in the mammary epithelium is highly coordinated yet distributed across phenotypically distinct epithelial subsets, giving rise to heterogeneous responses^8–10^.

Single-cell and spatial studies in normal breast and breast cancer have revealed that ERα and PR expression, and their downstream transcriptional programs, vary markedly across individual cells and lineages^11–13^. ERα⁺/PR⁺ and ERα⁺/PR⁻ populations coexist within the mammary epithelium and tumors, with only partial overlap between ERα and PR expression and limited correspondence between receptor positivity and proliferation^2,9,14^. In ERα⁺ breast cancer, variation in ER/PR expression and signaling output is increasingly recognized as a contributor to variable endocrine therapy responses and resistance^15–17^. These observations imply that hormone signaling output depends not only on receptor abundance but also on lineage identity, chromatin state, transcriptional co-regulators, and local microenvironmental cues. However, how these factors integrate to shape estrogen- and progesterone-responsive programs in intact epithelial systems remains poorly understood^2,18–21^.

Three-dimensional organoid models provide an attractive platform to address these questions because they preserve key aspects of mammary architecture, maintain basal and luminal lineage diversity, and better retain hormone receptor expression than traditional two-dimensional cultures^22–24^. Normal and tumor-derived breast organoids recapitulate ductal and acinar structures, sustain ER/PR expression for extended periods, and exhibit treatment responses that more closely mirror primary tissues^22,23,25^. However, even in organoids, ERα and PR expression often decline or shift over time, and most studies have not systematically examined hormone signaling at single-cell resolution across distinct mammary lineages or in relation to chromatin and co-regulator landscapes^23,26,27^.

Here, we investigated how hormone receptor abundance, transcriptional co-regulators, and cellular context shape estrogen and progesterone responses at single-cell resolution. To address this question, we used two complementary primary murine organoid systems. Organoids were generated from ERα enriched *Esr1*-*ZsGreen*⁺/PROM1⁺ luminal cells and from basal epithelial cells isolated from mouse mammary glands. Together, these systems provided complementary models to examine hormone responses in luminal cell-derived organoids and basal cell-derived organoids that acquire luminal-like identities in 3D culture. We found that estrogen and progesterone responses varied substantially across individual cells and were only weakly predicted by receptor transcript levels. Instead, variation in transcriptional co-regulators and microenvironmental cues are associated with to the magnitude and dynamics of hormone responses. Finally, by contrasting hormone response dynamics in basal cell-derived organoids with those in ERα⁺ breast cancer cells, we connect principles of hormone signaling heterogeneity in normal epithelium to those operating in disease. Together, these findings demonstrate that hormone signaling heterogeneity arises from multiple regulatory layers beyond receptor abundance and provide a framework for understanding variable endocrine responses in mammary epithelium and breast cancer.

## Results

### Mammary basal cells acquire luminal-like identities in organoid culture

To determine whether hormone receptor heterogeneity in the mammary gland reflects intrinsic cellular properties or arises from differences in microenvironmental cues, we employed 3D organoid cultures that allow precise control of the microenvironmental niche conditions. Organoids were generated from primary mouse mammary epithelium using two epithelial populations isolated by fluorescence-activated cell sorting. ERα+ luminal/progenitor cells were identified using the *Esr1*-*ZsGreen* knock-in reporter mouse line^28^ combined with PROM1 (prominin 1) staining, which enriches for an ERα+ luminal progenitor population responsible for luminal maintenance in vivo^29^. In parallel, basal epithelial cells were isolated from wild-type C57BL/6 mammary glands by excluding hematopoietic and endothelial lineages (CD31⁻ CD45⁻ TER119⁻) and gating on CD24⁺ CD29^hi^ cells^30^, a basal population with demonstrated mammary repopulating capacity^31^. These two sorted populations were used to directly compare their ability to generate hormone receptor-positive luminal cells in organoid culture (Fig. S1A). Despite their distinct cellular origins, both populations formed acinar-like organoids with similar morphologies, including cystic structures with intact lumens or budding architectures (Fig. S1B). Immunocytochemistry for ERα and PR revealed that receptor-positive nuclei occupied only a subset of cells within each organoid rather than being uniformly distributed, indicating that hormone receptor expression is heterogeneous in both ZsGreen⁺/PROM1⁺-derived and basal cell-derived organoids (Fig. 1A). This early observation suggested that only a fraction of cells in each system is directly hormone-competent, motivating further characterization of lineage composition and receptor landscapes before dissecting hormone-response heterogeneity.

**Figure 1.**
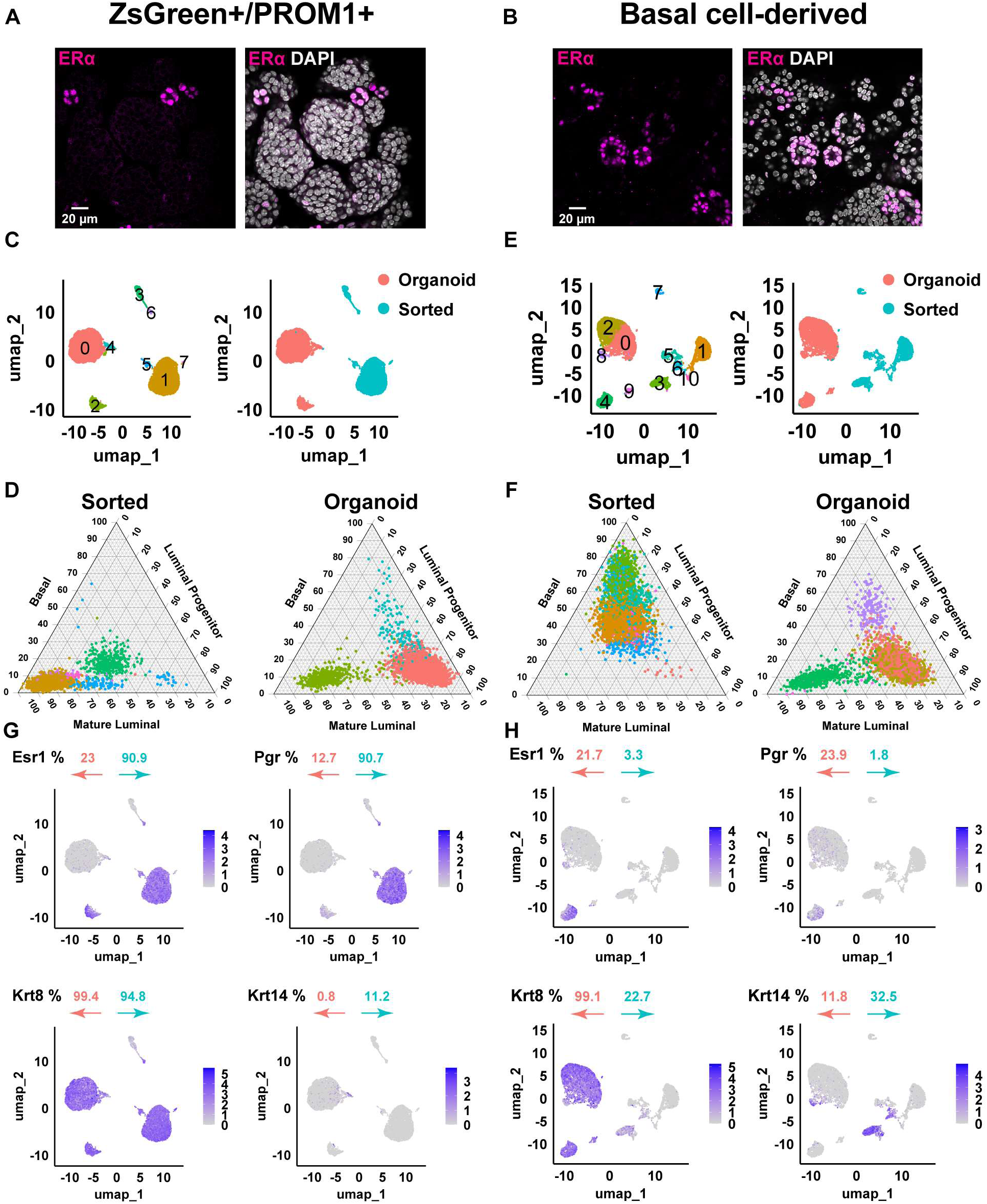
Lineage plasticity and restricted hormone receptor expression in mammary organoids (A) Representative immunofluorescence images of ERα (magenta) in ZsGreen⁺/PROM1⁺-derived organoids. Right panel shows ERα with DAPI nuclear counterstain. ERα-positive nuclei are detected in a subset of cells within each organoid. Scale bar, 20 µm. (B) Representative immunofluorescence images of ERα (magenta) in basal cell-derived organoids with DAPI nuclear counterstain. ERα expression is restricted to a minority of nuclei. Scale bar, 20 µm. (C) UMAP projections of single-cell RNA-seq profiles from ZsGreen⁺/PROM1⁺-sorted cells and their derived organoids. Left, unsupervised clusters; right, cells colored by origin (sorted versus organoid), showing distinct transcriptional states following organoid culture. (D) UMAP projections of basal-sorted cells and basal cell-derived organoids. Left, cluster annotation; right, cells colored by origin. Sorted and organoid populations occupy non-overlapping transcriptional spaces. (E) Ternary plot showing lineage gene-set scores (basal, luminal progenitor, mature luminal) for ZsGreen⁺/PROM1⁺-sorted cells and derived organoids. Organoids shift toward luminal progenitor-like identities relative to sorted cells. (F) Ternary plot of lineage scores for basal-sorted cells and basal cell-derived organoids. Basal-sorted cells score predominantly as basal, whereas basal cell-derived organoids acquire luminal progenitor and mature luminal signatures. (G) UMAP feature plots showing expression of Esr1, Pgr, Krt8, and Krt14 in ZsGreen⁺/PROM1⁺-sorted cells and corresponding organoids. Percentages indicate the fraction of gene-positive cells in sorted (blue) and organoid (orange) samples. (H) UMAP feature plots showing expression of Esr1, Pgr, Krt8, and Krt14 in basal-sorted cells and basal cell-derived organoids. Basal cell-derived organoids lack a basal transcriptomic signature and exhibit restricted hormone receptor expression within luminal-like populations.

To determine whether mammary epithelial cells maintain or alter their transcriptional states in 3D culture, we performed scRNA-seq on sorted ERα^+^ luminal and basal populations and on their corresponding organoids. Unsupervised clustering of 10x Genomics scRNA-seq data revealed five transcriptionally distinct clusters among the ERα⁺ sorted luminal cells and three clusters in their derived organoids (Fig. 1C). When visualized in UMAP space and colored by sample origin, sorted and organoid cells formed completely non-overlapping clusters, indicating a global and culture-driven shift in transcriptional state following organoid formation (Fig. 1C).

Lineage identities were assigned using a gene set-scoring approach based on publicly curated signatures for basal, luminal progenitor, and mature luminal lineages^32,33^. In a ternary plot of ZsGreen^+^/PROM1^+^ luminal samples, the largest sorted cluster scored predominantly for mature luminal identity, with smaller clusters enriched for progenitor signatures (Fig. 1D). By contrast, the dominant organoid cluster shifted toward a progenitor-like profile, while smaller subsets retained mature luminal characteristics (Fig. 1D; Table S1). Although PROM1 was used to identify luminal progenitors in vivo, our ternary scoring revealed unexpected enrichment for fully differentiated luminal signatures rather than progenitors. These analyses indicate that ZsGreen^+^/PROM1^+^ luminal cells are predominantly mature at the time of sorting, whereas organoid cultures contain a higher proportion of progenitor-like states. This change in composition could reflect both partial reprogramming of mature luminal cells or preferential expansion of a minor progenitor-enriched subpopulation captured by the PROM1 gate. Organoid cultures also displayed an increased fraction of cycling cells with S-phase enrichment relative to sorted samples (Fig. S1C; Table S1), indicating a redistribution of cell-cycle states alongside the shift toward progenitor-like identities.

Applying the same workflow to basal-sorted and basal-derived organoid cells revealed six distinct clusters in the basal-sorted population (including one representing endothelial contamination, Cluster 1) and five clusters in the organoids (Fig. 1E). Endothelial contamination (*Cd31⁺* clusters [Pecam1^+^]; Fig. S1D) persisted despite using a combined exclusion channel (“lineage cocktail”) to gate out hematopoietic and endothelial contaminants, and we retained them to preserve global heterogeneity, although they do not survive long-term 3D culture. Gene-set scoring and ternary plot analysis showed that basal-sorted cells formed a dominant basal cluster consistent with their lineage identity. In contrast, basal cell-derived organoid clusters largely lost this basal signature and instead acquired luminal progenitor transcriptional profiles, with a minority also exhibiting mature luminal scores (Fig. 1F; Table S1). Basal cell-derived organoids also showed a marked shift in cell-cycle state, with an increased proportion of G1-phase cells and reduced S and G2/M fractions relative to basal-sorted cells (Fig. S1E; Table S1), suggesting a shift toward G1 enrichment during basal-to-luminal conversion.

Analysis of luminal and basal lineage scores across all cells showed that ZsGreen^+^/PROM1^+^ organoids retained luminal lineage features, whereas basal cell-derived organoids exhibited a marked reduction in basal transcriptomic scoring by our criteria. Instead, basal cell-derived cultures were dominated by luminal progenitor and mature luminal-like populations in 3D culture (Fig. 1E-F; Table S1). These findings were consistent with substantial basal-to-luminal plasticity under organoid conditions, although we cannot exclude contributions from selective survival or expansion of rare luminal-like cells present at the time of sorting.

To define the baseline hormone receptor landscape in each organoid system prior to functional hormone stimulation, we quantified *Esr1* and *Pgr* expression, while *Krt8* (keratin 8) and *Krt14* (keratin 14) were used to delineate lineage identity. In both organoid models, roughly one-fifth of cells were ERα^+^. Notably, basal cell-derived organoids contained nearly twice the fraction of PR^+^ cells compared with ZsGreen⁺/PROM1⁺ organoids (∼ 24% vs ∼ 13%) (Fig. 1G & H; Fig. S1F; Table S1). *Krt8* and *Krt14*, canonical luminal and basal markers, respectively, indicated a predominantly luminal composition across organoid conditions. Thus, hormone receptor-positive cells resided within a largely luminal compartment in both systems. Overall, these analyses show that both organoid systems are predominantly luminal in composition yet contain restricted and quantitatively distinct hormone receptor-positive subpopulations. The differential distribution of *Pgr*^+^ cells, together with the sparse *Esr1*^+^ compartment, defines distinct baseline receptor landscapes that set the stage for interrogating hormone-response heterogeneity.

### Basal organoid cells acquire lineage specific markers and luminal-like chromatin features in 3D culture

To further define lineage-associated molecular features across sorted and organoid populations, we examined expression of hormone receptor and luminal gene signatures (Fig. 2A & B). *Esr1* and *Pgr* expression delineated hormone-competent luminal compartments in both systems (Fig. 1G & H). Within these receptor-positive populations, mature luminal markers including *Foxa1* and *Ly6a* (*Sca1*) were co-expressed, consistent with a differentiated hormone-sensing identity. *Foxa1*, a pioneer factor required for ERα chromatin engagement^34^, and *Sca1*, a surface marker enriched in mature luminal cells, further define these compartments ^35^. The luminal regulator *Gata3*, a key pioneering factor required for luminal lineage specification and sustained ERα signaling^36,37^, was broadly expressed across both progenitor-like and mature luminal populations, while *Prlr*, a hallmark of hormone-responsive luminal cells^38,39^, was also detected within luminal clusters of both models. Notably, *Prlr* expression decreased in ZsGreen⁺/PROM1⁺ organoids relative to sorted cells, whereas it increased in basal cell-derived organoids compared with their sorted counterparts, revealing reciprocal modulation of mature luminal features across systems.

**Figure 2.**
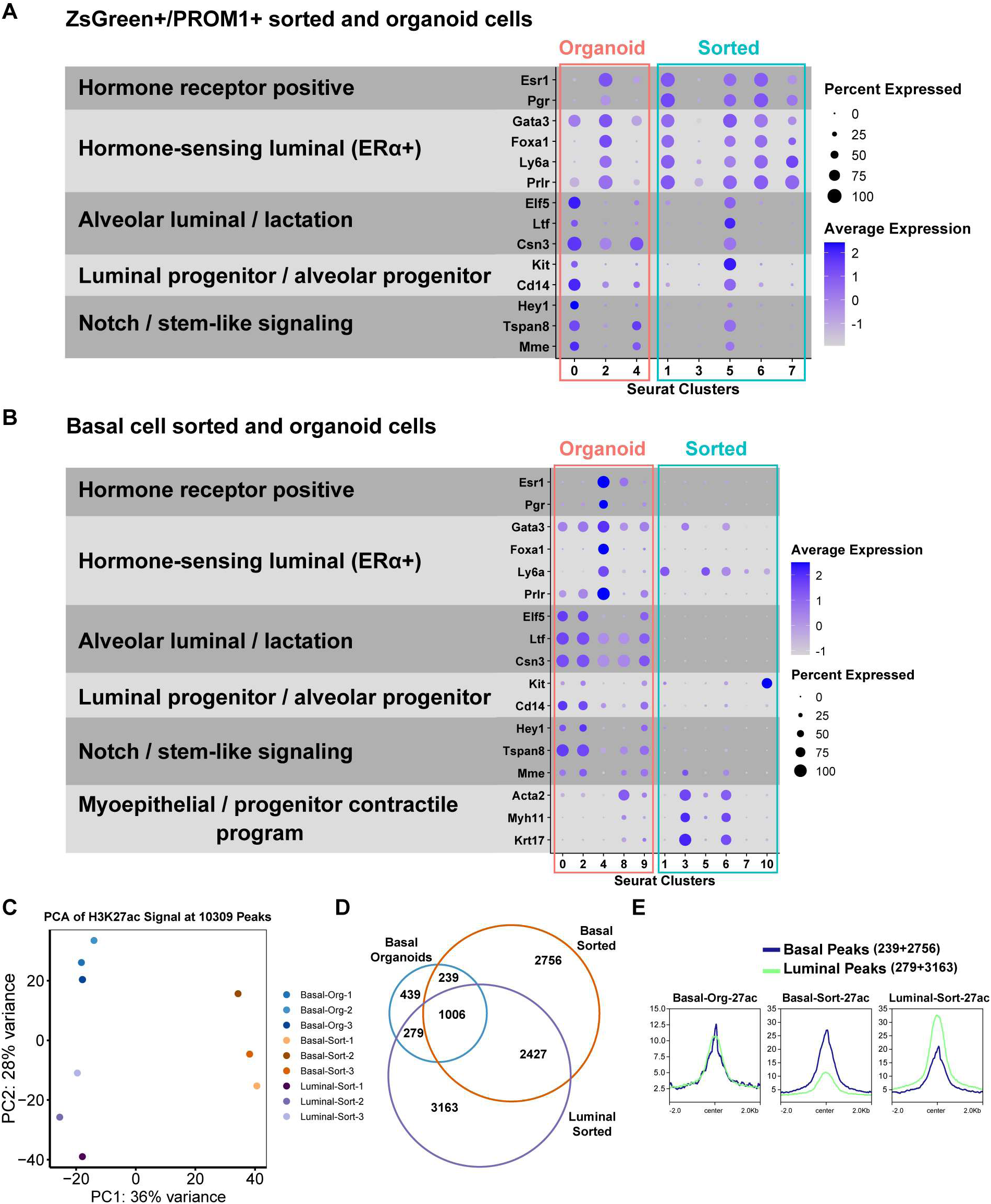
Organoid culture induces a hybrid luminal regulatory state in basal cells (A) Dot plot of lineage-associated gene expression across clusters from ZsGreen⁺/PROM1⁺-sorted cells and derived organoids. Genes are grouped by functional categories. Dot size represents the percentage of expressing cells and color indicates average scaled expression. (B) Dot plot of lineage-associated gene expression across clusters from basal-sorted cells and basal cell-derived organoids. (C) Principal component analysis of H3K27ac CUT&Tag profiles from basal-sorted, luminal-sorted, and basal cell-derived organoid samples. (D) Venn diagram showing overlap of H3K27ac peaks among basal-sorted, luminal sorted, and basal cell-derived organoid samples. (E) Aggregate H3K27ac signal profiles at basal-associated and luminal-associated enhancer regions across the indicated conditions.

*Elf5*, *Ltf*, and *Csn3* were prominently expressed in luminal progenitor populations identified by ternary plot analysis in both ZsGreen⁺/PROM1⁺ and basal cell-derived systems (Fig. 2A, B). These genes are established markers of alveolar progenitor-like states associated with alveologenesis and luminal fate specification. *Elf5* functions as a master regulator of alveolar differentiation^40–42^, while *Ltf*^11,43^ and *Csn3*^44^ further reinforce this transcriptional program. Additional progenitor-associated markers, including *Hey1*^45^, *Tspan8*^46^, *Mme*^47^, *Kit*, and *Cd14*, were similarly enriched within these luminal progenitor compartments in both systems, consistent with their reported roles in progenitor maintenance and colony-forming capacity^32,48–50^. Together, these patterns support the presence of an alveolar progenitor-like transcriptional state within organoid cultures, including a subset of basal cell-derived cells.

Conversely, basal lineage-associated genes including *Acta2*, *Myh11*, *Sox10*, and *Krt17* were expressed almost exclusively in basal-sorted cells^51–53^ (Fig. 2B). *Sox10* expression was also detected in a subset of ZsGreen⁺/PROM1⁺ organoid cells (Fig. 2A), consistent with the presence of progenitor-associated features within this population. In parallel, basal cell-derived organoid cells expressed mature luminal markers, further illustrating the redistribution of lineage-associated transcriptional programs in 3D culture. Together, these patterns define the molecular distinctions between progenitor-like and mature hormone-sensing states across systems and clarify the lineage context in which hormone receptor heterogeneity is examined.

To investigate whether transcriptional reprogramming in basal cell-derived organoid cells is accompanied by epigenetic remodeling, we profiled enhancer-associated chromatin states using Cleavage Under Targets and Tagmentation (CUT&Tag)^54^ for the active enhancer mark H3K27ac. Because basal cell-derived organoids undergo a pronounced basal-to-luminal lineage transition, we focused these analyses on basal-sorted epithelial cells and their derived organoids, using freshly isolated luminal epithelial cells as a reference endpoint for luminal enhancer states. This design enabled direct comparison of lineage-specific enhancer landscapes across basal, organoid, and luminal epithelial populations. In addition, we profiled the repressive histone mark H3K27me3 in parallel, which is summarized in the form of metadata visualization and heatmap. These datasets allowed us to assess whether chromatin accessibility dynamics in basal organoids reflect the luminal lineage transition observed in our scRNA-seq analysis.

Principal component analysis of H3K27ac profiles revealed clear separation between basal-sorted and luminal-sorted samples, consistent with their distinct enhancer usage and underlying transcriptional programs (Fig. 2C). In contrast, basal cell-derived organoid samples no longer clustered with basal-sorted cells and instead shifted toward the luminal-sorted samples along the principal components. This shift indicates that basal organoid cells undergo substantial chromatin reorganization during 3D culture, acquiring enhancer features that partially converge toward a luminal epigenetic state. However, they remain distinct from fully differentiated luminal cells. These results are in line with the transcriptional evidence showing partial adoption of luminal identity by basal cell-derived organoid cells. Given that CUT&Tag is a bulk method, this intermediate profile likely reflects the cellular heterogeneity within the organoids.

Consistent with this observation, Venn diagram analysis of H3K27ac peaks demonstrated both shared and unique enhancer elements across the three conditions (Fig. 2D). A total of 6707 peaks were identified in basal-sorted cells, 7114 peaks in luminal-sorted cells, and 1963 peaks in basal cell-derived organoids. The reduced number of peaks detected in organoid samples likely reflects lower CUT&Tag efficiency in 3D conditions. However, biological differences in chromatin accessibility potentially induced by basement membrane-associated signaling cues may also contribute. Notably, basal cell-derived organoids contained a distinct set of 439 unique H3K27ac peaks not detected in either sorted population, suggesting the emergence of organoid-specific regulatory elements.

Importantly, basal organoid cells shared comparable numbers of enhancer peaks with both basal-sorted (239 peaks) and luminal-sorted (279 peaks) populations. This balanced overlap supports the notion that basal cell-derived organoids are no longer epigenetically aligned with their cell of origin but instead adopt an intermediate, hybrid enhancer landscape indicative of lineage transition. To further contextualize these changes, we classified H3K27ac peaks as basal- or luminal-associated based on their presence in the corresponding sorted populations. Metadata-based visualization revealed that basal cell-derived organoids harbor both basal and luminal enhancer signatures at similar relative levels, whereas basal-sorted and luminal-sorted cells showed the expected enrichment for their respective lineage-associated enhancer peaks (Fig. 2E and Fig. S2A). When we overlaid H3K27me3 signal onto these H3K27ac-defined regions, basal cell-derived organoids showed higher H3K27me3 at basal-associated peaks than at luminal-associated peaks (Fig. S2A). In basal-sorted cells, luminal-associated peaks carried higher H3K27me3 than basal-associated peaks, whereas in luminal-sorted cells, basal-associated peaks were more heavily marked than luminal-associated peaks. Collectively, these patterns are consistent with lineage-specific repression across the shared enhancer space. Genome browser views of representative luminal-associated loci further illustrate that basal cell-derived organoids acquire H3K27ac enrichment comparable to luminal-sorted cells but not basal-sorted cells (Fig. S2B). Together with our scRNA-seq data, these epigenetic analyses demonstrate that basal cells undergoing organoid culture not only alter their gene expression programs but also acquire luminal-like enhancer and repression features, providing independent evidence of a chromatin-level transition toward a luminal regulatory state.

### Basal cell-derived mammary organoids exhibit robust and extended hormone responsiveness in 3D culture

Having established the transcriptional identities of ZsGreen⁺/PROM1⁺ and basal cell-derived organoids, we next investigated whether these systems maintain functional hormone responsiveness in 3D culture. To evaluate hormone signaling in these systems, organoids were treated with 100 nM E2, 1 µM P4, E2+P4, or vehicle control for 4h and 24h. Bulk RNA-seq of E2-treated ZsGreen⁺/PROM1⁺ organoids compared with vehicle controls revealed hormone-responsive genes that passed statistical thresholds, but the vast majority exhibited low fold-changes, indicating that the overall transcriptional response was weak (Fig. 3A and Fig. S3A). Because the sorted ZsGreen⁺/PROM1⁺ population is enriched for hormone-sensing luminal cells, we compared freshly sorted cells with their derived organoids. This analysis revealed a marked transcriptomic shift upon adaptation to 3D culture, indicating that gene expression programs were extensively reorganized during culture (Fig. 3B). Consistent with this global shift, sorted cells exhibited high expression of *Esr1*, *Pgr*, and *Foxa1*, key markers of hormone-sensing luminal cells. In contrast, organoid cultures showed reduced expression of these markers and elevated levels of luminal progenitor-associated genes *Kit*, *Cd61* (*Itgb3*), *Cd49b* (*Itga2*) (Fig. S3B). *Esr1* transcripts were not readily detected in bulk RNA-seq of organoid cultures, likely because rare ERα+ cells visualized by immunostaining in Fig. 1A become masked in averaged transcript measurements. Similarly, PR protein was detectable only in a small subset of cells in ZsGreen⁺/PROM1⁺ organoids (Fig. S3C), supporting the idea that hormone receptor levels vary from cell to cell within organoid culture. Together with low ERα/PR protein detection, these data support a marked reduction in ERα+ mature luminal cells following adaptation to 3D culture. This reduction aligns with the transcriptional shift observed by scRNA-seq, consistent with organoid cultures being enriched for luminal progenitor-like cells (Fig. 1E).

**Figure 3.**
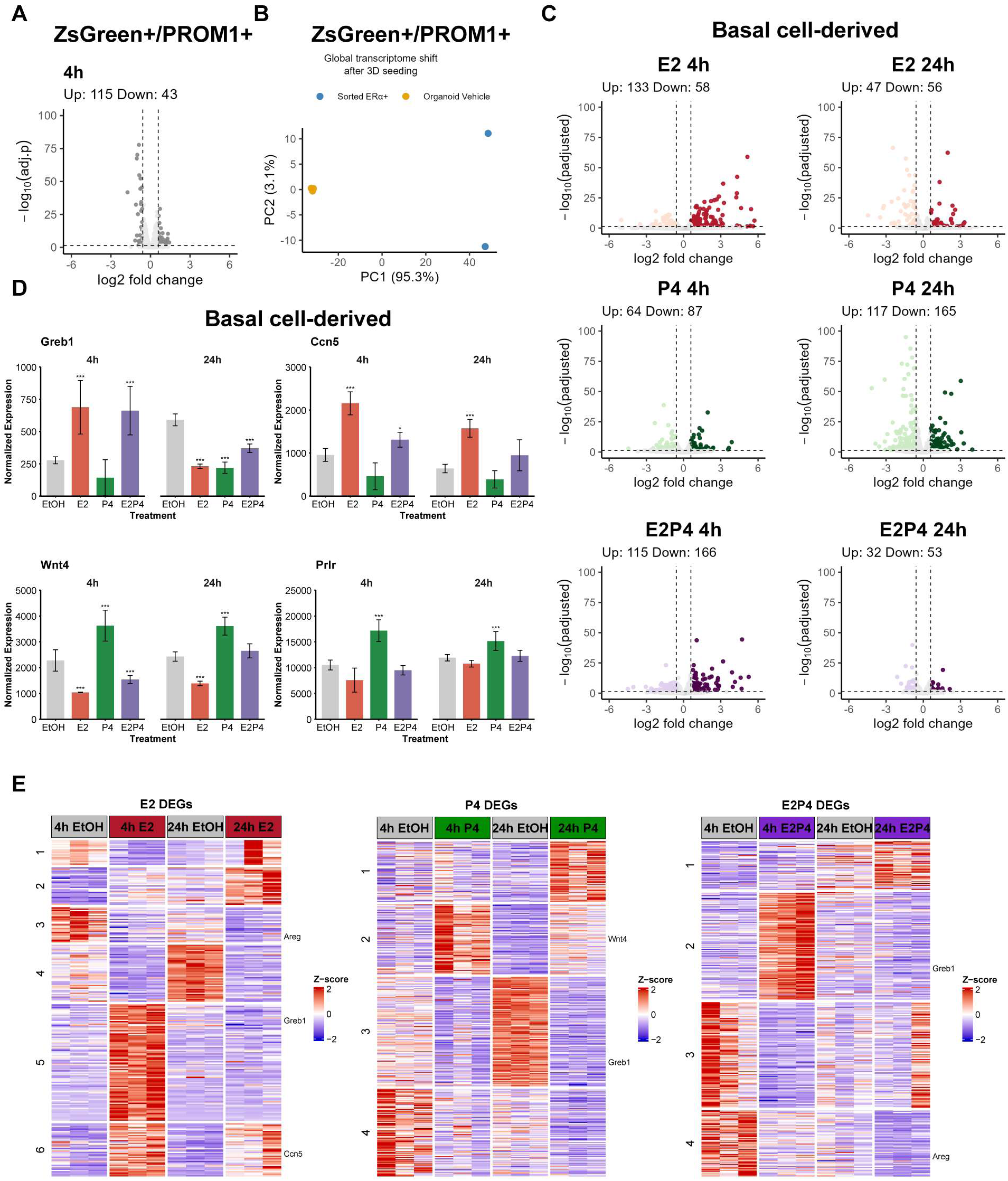
Functional estrogen and progesterone signaling is preserved in basal organoids (A) Volcano plot of differentially expressed genes (DEGs) in ZsGreen⁺/PROM1⁺ organoids following E2 treatment. DEGs were defined as adjusted p value < 0.05 and |log2 fold change| ≥ 0.585. Although statistically significant genes were detected, most exhibited low fold changes. (B) Principal component analysis comparing freshly sorted ZsGreen⁺/PROM1⁺ cells and their derived organoids, demonstrating global transcriptional reorganization upon adaptation to 3D culture. (C) Volcano plots of DEGs in basal cell-derived organoids following treatment with E2, P4, or E2+P4 for 4h and 24h. Significant transcriptional responses are observed at both early and late time points. (D) Normalized expression of canonical estrogen targets (*Greb1*, *Ccn5*) and progesterone targets (*Wnt4*, *Prlr*) across hormone treatments at 4h and 24h in basal cell-derived organoids. Bars represent mean ± SE across biological replicates (n = 3 per condition). Statistical significance was determined using adjusted p-values from DESeq2 differential expression analysis (Wald test with Benjamini-Hochberg correction). Asterisks denote adjusted p-values (*padj < 0.05, **padj < 0.01, ***padj < 0.001). (E) Heatmaps of DEGs for E2, P4, and E2+P4 treatments at 4h and 24h in basal cell-derived organoids. Hierarchical clustering identifies six temporal clusters in the E2 condition and four clusters each in the P4 and E2+P4 conditions, reflecting distinct hormone-specific expression programs.

In contrast, basal cell-derived organoids remained highly responsive to hormonal stimulation. Volcano plots revealed a limited number of statistically significant differentially expressed genes (DEGs) in response to hormone conditions at both 4h and 24h (Fig. 3C; Table S2). Because *Esr1*-expressing cells represent only a minority of the organoid population (∼ 20% by scRNA-seq; Fig. 1), hormone-induced transcriptional responses are diluted in bulk RNA measurements, such that only the most strongly regulated targets are detected as DEGs. To further illustrate hormone-responsiveness, we examined canonical E2 targets growth regulation by estrogen in breast cancer 1 (*Greb1*) and cellular communication network factor 5 (*Ccn5*), as well as P4 targets Wnt family member 4 (*Wnt4*) and prolactin receptor (*Prlr*) from the RNA-seq data. *Greb1* was induced at 4h by E2 and E2+P4, but not by P4 alone, and showed reduced expression at 24h across hormone treatments due to higher baseline levels in vehicle (EtOH) control. *Ccn5* was also upregulated at 4h in E2 and E2+P4 conditions, while at 24h it remained elevated only with E2, with no changes under P4 or combined treatment. The progesterone target *Wnt4* was selectively induced by P4 at both 4h and 24h, whereas it was downregulated by E2 and E2+P4 at 4h and suppressed by E2 alone at 24h. *Prlr* displayed a consistent progesterone-driven pattern, with strong induction at both 4h and 24h under P4 treatment, and no significant changes in other conditions. Together, these data demonstrate intact estrogen and progesterone signaling, reflected by both global transcriptomic profiles and the differential regulation of well-established hormone targets (Fig. 3D). Notably, this endocrine responsiveness was preserved in long-term culture, with organoids remaining hormone-responsive after approximately 10 months of continuous culture spanning seven serial passages. Thus, under matched 3D culture conditions, basal cell-derived organoids retain functional hormone signaling capacity, whereas ZsGreen⁺/PROM1⁺ organoids exhibit markedly attenuated transcriptional responses to E2 treatment, indicating that luminal lineage identity alone is not sufficient to predict estrogen responsiveness in this system.

To capture global expression dynamics over time, we generated heatmaps of DEGs for each hormone treatment at 4h and 24h. Hierarchical clustering revealed distinct temporal expression patterns, with six major clusters identified in the E2 datasets, four clusters in the P4 datasets, and four clusters in the combined E2+P4 datasets, highlighting diverse temporal patterns of hormone regulation (Fig. 3E). In the E2 condition, six clusters encompassed several distinct behaviors. Cluster 1 contained genes that were modestly elevated in 4h EtOH controls but suppressed by E2 at the early (4h) time point, with only partial recovery by 24h. Cluster 2 comprised genes with delayed E2-specific induction, becoming strongly activated only at 24h. Clusters 3 and 4 showed sustained E2-driven repression across both time points. Cluster 5 represented rapid but transient early-response genes induced at 4h but returning to baseline by 24h, whereas Cluster 6 displayed sustained induction, with transcripts elevated at 4h and remaining increased at 24h.

By contrast, P4-treated basal organoids resolved into only four clusters, representing more streamlined transcriptional trajectories. These included genes with strong late induction, subsets transiently induced at 4h and reduced by 24h, and others suppressed across both time points. The E2+P4 condition also produced four clusters, characterized by combined effects such as reinforced repression or attenuation of E2-driven induction, reflecting points of synergy and antagonism between estrogen and progesterone signals. Together, these analyses highlight that E2 elicits the most complex induction kinetics and dynamic temporal responses, whereas P4 and combined hormone treatments generate fewer, more coordinated expression programs.

### Heterogeneous single-cell hormone responsiveness correlates with co-regulator abundance and differs across cellular context

To determine whether normal hormone-responsive cells exhibit heterogeneous responses even when microenvironmental conditions are controlled, we performed scRNA sequencing on basal cell-derived organoid cells treated with 10 nM E2 for 4h and 16h or 1 µM P4 for 4h, focusing our analysis on the ERα^+^ subpopulation. We used 10 nM E2 for scRNA-seq because the single-cell platform provides sufficient sensitivity to detect transcriptional responses at physiological hormone levels, even when ERα^+^ cells represents only a subset of the culture (Fig. S4A). This approach contrasted with the bulk RNA-seq experiments, where we used 100 nM E2 to ensure that responses from the ∼ 20% ERα^+^ cells were not diluted by the larger ERα^-^ population in population-averaged measurements. By focusing our analysis on the ERα^+^ subpopulation, we detected hormone-responsive genes with substantially greater sensitivity than bulk RNA-seq, with hundreds of differentially expressed genes identified for each hormone condition (Fig. 4A; Table S3).

**Figure 4.**
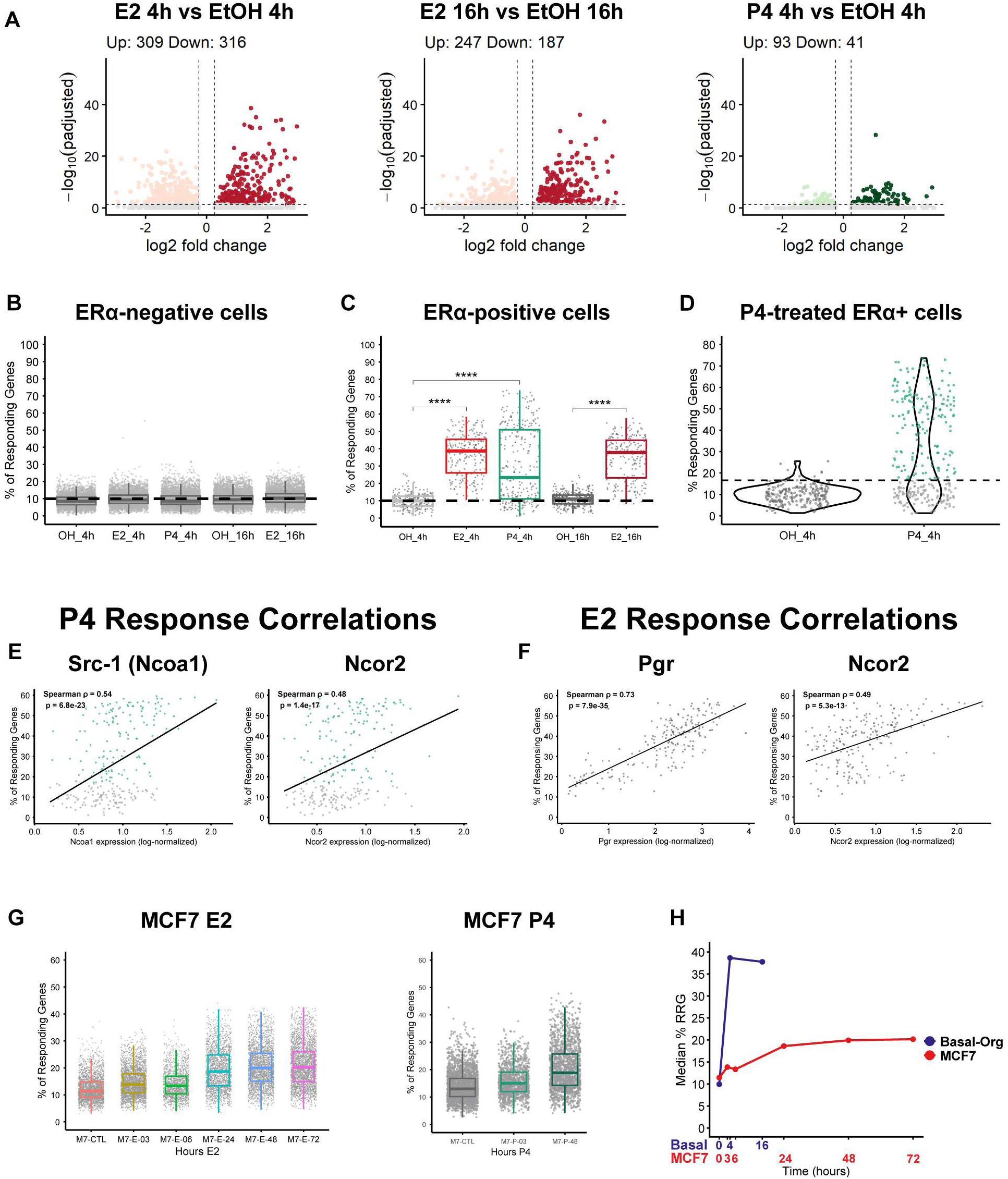
Heterogeneous hormone responses and distinct temporal kinetics in organoids and MCF7 cells (A) Volcano plots showing differentially expressed genes (DEGs) in ERα^+^ basal cell-derived organoid cells treated with E2 pr P4 for 4h and 16h relative to vehicle control. DEGs were defined as adjusted p value < 0.05 and |lo2 fold change| ≥ 0.25. (B) Percentage of responding genes (%RRG) per cell in ERα^-^ populations across treatments and time points. (C) %RRG per cell in ERα^+^ populations following E2 or P4 treatment for 4h and 16h. (D) Distribution of %RRG values on ERα^+^ cells following P4 treatment at 4h. (E) Correlation between co-regulator expression (Ncoa1, Ncor1, Ncor2) and P4-induced %RRG in ERα^+^ cells. Spearman correlation coefficients are indicated. (F) %RRG in published single-cell RNA-seq datasets of MCF7 cells treated with E2 or P4 at indicated time points. (G) Median %RRG over time in basal cell-derived organoids and MCF7 cells.

To further characterize cell-to-cell variability in hormone responses, we asked how many hormone-responsive genes showed a detectable change in each cell. To calculate the “ratio of responding genes” (%RRG), we first determined the mean and variability of each hormone-responsive upregulated DEG in vehicle-treated cells. We then computed, for each treated cell, the fraction of these genes that were significantly upregulated, as previously described^55^. We calculated the percentage of responding genes detected per cell separately in ERα-negative (ERα^-^) and ERα-positive (ERα^+^) populations. ERα^-^ cells served as an internal negative control, exhibiting a stable baseline proportion of responding genes across all treatments and time points, with median values clustered around 9% (Fig. 4B; Table S3). Having established the stable baseline in ERα^-^ cells, we next quantified hormone-induced transcriptional dynamics within the ERα^+^ compartment. ERα^+^ cells exhibited robust and time-dependent responses: the median %RRG increased roughly fourfold after 4h E2 treatment compared with vehicle, indicating that, on average, each ERα^+^ cell upregulated nearly one-third of all hormone-responsive genes (Fig. 4C; Table S3). P4 exposure produced an intermediate increase in %RRG, and at 16h the E2-treated ERα^+^ cells maintained elevated %RRG relative to controls, consistent with rapid onset and sustained transcriptional activation in the ERα^+^ compartment (Fig. 4C; Table S3). Overall, E2-induced transcriptional responses in basal cell-derived organoids demonstrate rapid kinetics, rising within 4 hours and plateauing through 16 hours.

Notably, P4-treated samples displayed bimodal distribution of %RRG values, with one subpopulation maintaining near-baseline values and another exhibiting markedly elevated %RRG values between 20-80% (Fig. 4D; Fig. S4B). Because transcriptional responses to hormones are often influenced by receptor levels, we first asked whether *Pgr* transcript abundance correlated with the magnitude of P4-induced %RRG across ERα⁺ cells. Although *Pgr* expression is detectable in untreated basal cell-derived organoids (Fig. 1H), *Pgr* expression showed only a weak and inconsistent association with P4 responsiveness (Pearson r = -0.16; Spearman ρ = -0.12), suggesting that *Pgr* transcript abundance alone does not account for the observed heterogeneity, consistent with the possibility that PR protein levels are not strictly determined by mRNA abundance (Fig. S4C; Table S3). Because transcriptional co-regulators modulate nuclear receptor activity, we asked whether variability in key co-regulators (*Ncoa1*, *Ncor1*, *Ncor2*) contributes to the cell-to-cell heterogeneity in hormone responses. Correlation analyses revealed that, among the co-regulators tested, *Ncoa1* (*Src-1*) (ρ = 0.54) and *Ncor2* (ρ = 0.48) expression levels were significantly associated with P4-induced %RRG (Fig. 4E; Table S3), whereas Ncor1 and Ncoa3 showed no correlation (Fig. S4C; Table S3). To determine whether similar relationships exist for estrogen signaling, we performed the same analysis using E2-induced responsive genes. In this context, *Pgr* expression showed the strongest correlation with E2-induced %RRG (Spearman ρ = 0.73), consistent with its role as a canonical ERα target gene. Notably, *Ncor2* expression also exhibited a positive association (Spearman ρ = 0.49) with E2 response magnitude (Fig. 4F; Table S3) while *Esr1*, *Ncor1*, and *Scr-1* displayed only weak or non-significant relationships (Fig. S4D; Table S3). Together, these results suggest that variability in receptor and co-regulator expression is significantly associated with to the heterogeneous magnitude of acute hormone responses across ERα⁺ cells.

To contrast these rapid hormone-response dynamics in basal cell-derived organoids with ERα+ breast cancer cells, we analyzed published scRNA-seq profiles of E2- or P4-treated MCF7 cells, generated by Ors et al.^56^. In MCF7 cells, the %RRG remained largely unchanged during the early time points, indicating minimal acute transcriptional activation, and increased by approximately 50% only at later stages following prolonged hormone exposure (Fig. 4G; Table S3). P4 treatment displayed a similar pattern with negligible changes at 3h (12.1%) followed by an increase at 48h (16.4%). Together, these findings reveal that hormone-induced transcriptional responses follow markedly different temporal trajectories in basal organoids versus MCF7 cells. Basal organoids show rapid activation, reaching a high plateau of ∼38% responding genes by 4-16h, whereas MCF7 cells display minimal early responses and only gradually approach a lower plateau of ∼20% over 24-72h (Fig. 4H). These contrasting kinetics highlight how cellular context influences the timing and magnitude of hormone-driven transcriptional programs, indicating that estrogen responses differ between normal mouse basal cell-derived organoids and human ERα+ breast cancer cells. Nevertheless, both systems retain substantial cell-to-cell variability in the proportion of responding genes, indicating that heterogeneous endocrine signaling is a conserved feature of estrogen-responsive epithelial cells in these culture models.

### Hormone receptor expression is dynamically remodeled by hormonal and microenvironmental cues in basal cell-derived mammary organoids

To establish the quantitative baseline of hormone receptor expression at the organoid level, we measured ERα and PR protein across individual basal cell-derived organoids. ERα and PR immunocytochemistry was performed in basal cell-derived organoids, and 20 organoids were analyzed for protein quantification. Confocal z-stacks spanning the full thickness of each organoid were acquired, and 3D reconstructions were used to quantify the percentage of ERα+ and PR+ cells per organoid. Representative images of ERα and PR staining are shown, with K8 (keratin 8) used as a luminal marker, given that basal cell-derived organoids adopt luminal-like identities in 3D culture (Fig. 5A).

**Figure 5.**
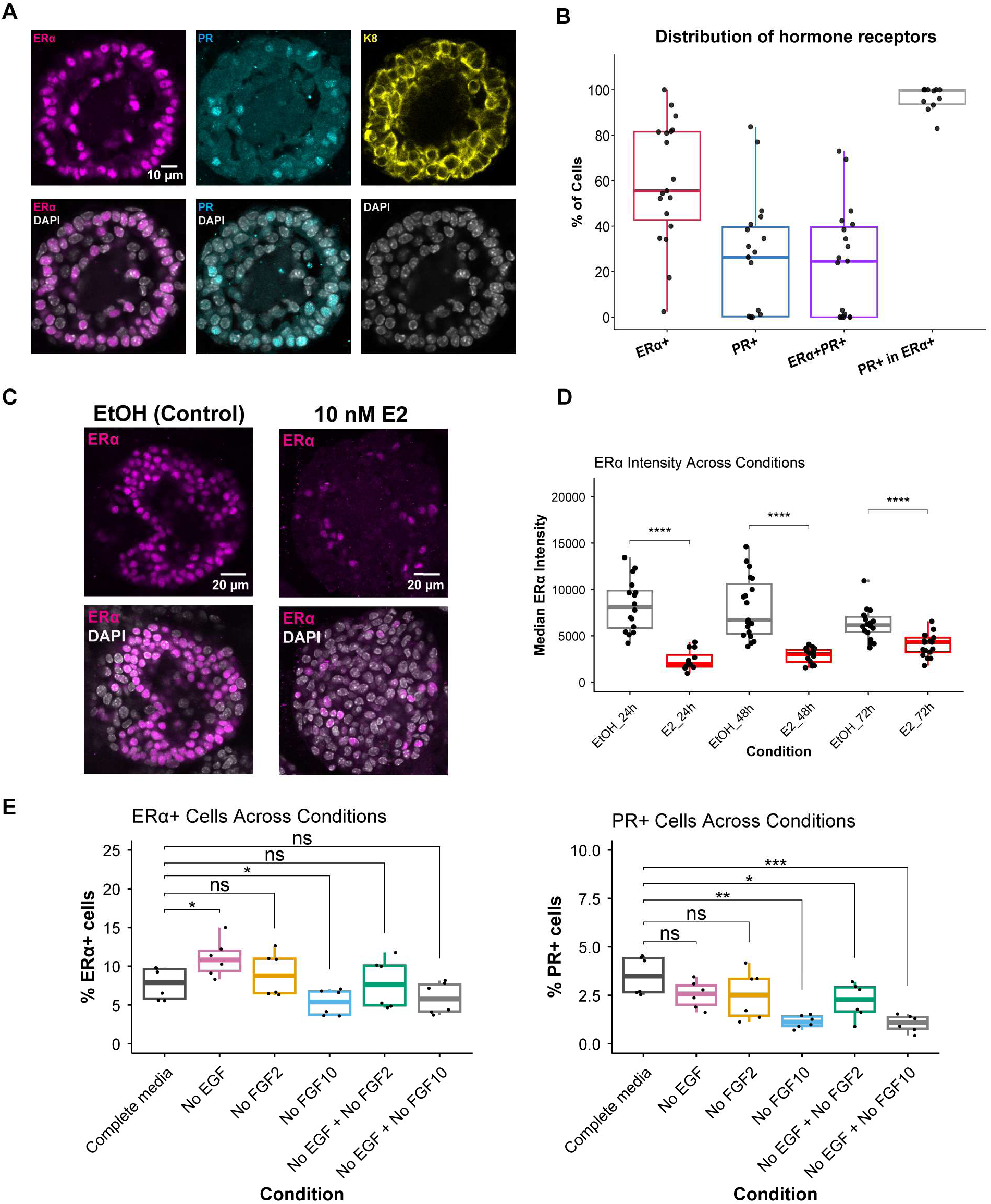
Context-dependent regulation of ERα and PR in mammary organoids (A) Representative confocal images of ERα (magenta), PR (cyan), and K8 (yellow) immunostaining in basal cell-derived organoids. DAPI marks nuclei (gray). Scale bar, 10 µm. (B) Quantification of ERα^+^, PR^+^, and ERα^+^/PR^+^ cell fractions across individual organoids (n = 20). Each point represents one organoid. (C) Representative images of ERα immunostaining in organoids treated with vehicle (EtOH) or 10 nM E2 for 48h. Scale bar, 20 µm. (D) Quantification of median ERα fluorescence intensity following 10 nM E2 treatment for 24h, 48h, and 72h compared with vehicle controls. (E) Percentage of ERα^+^ cells (left) and PR^+^ cells (right) under indicated growth factor conditions. Tile-scan imaging was used to quantify receptor-positive cells across entire wells. Statistical comparisons are indicated.

Quantification of ERα+ cells across individual organoids revealed marked heterogeneity (Fig. 5B). While some organoids contained predominantly ERα-expressing cells, others were largely ERα-negative, with a median of 55.6% ERα+ cells across organoids. PR protein levels were lower overall but similarly heterogeneous, with a median of 26.4% PR+ cells and values ranging from absent to nearly 80% per organoid. The median fraction of ERα+/PR+ cells was 24.6%, and 99.8% of PR+ cells also expressed ERα, indicating that PR is largely restricted to the ERα+ compartment, whereas a subset of ERα+ cells lacks PR expression.

We next tested whether ERα protein levels in basal cell-derived organoids are dynamically regulated by estrogen signaling. Estrogen binding to ERα is known to trigger transcriptional activation followed by proteasome-mediated receptor degradation^3,57,58^. We therefore asked whether basal cell-derived organoids recapitulate this canonical ERα biology. Organoids were treated with 10 nM E2 for 24h, 48h, or 72h, followed by immunostaining. Representative images from 48h comparing vehicle (EtOH) control- and E2-treated samples show a pronounced reduction in ERα-expressing cells upon E2 treatment (Fig. 5C). Quantitative analysis of ERα signal intensity demonstrated a significant decrease in ERα protein levels in E2-treated samples relative to controls at all time points (Fig. 5D). Interestingly, temporal analysis revealed opposing trends in ERα levels between control and E2-treated samples. In control organoids, ERα levels progressively declined over time, whereas ERα levels increased over time in E2-treated samples. Although ERα levels remained significantly different between control and E2 conditions at 72h, these opposing trajectories resulted in partial convergence of ERα protein levels (Fig. S5A). These opposing trajectories suggest that ligand-dependent feedback helps maintain ERα expression over time in E2-treated organoids, whereas in the absence of estrogen ERα levels progressively decline.

We then asked whether ERα and PR protein levels could be modulated by altering the organoid microenvironment independently of hormone treatment. Based on prior studies demonstrating that FGF signaling influences mammary epithelial cell state in organoids^59,60^, we tested the effects of FGF2 and FGF10 withdrawal. In addition, given the well-established crosstalk between EGF and ERα signaling pathways^61,62^, we also examined the effect of EGF withdrawal. To quantify receptor expression changes, tile-scan images were acquired to capture all organoids within a well, minimizing sampling bias (Fig. S5B). Multiple focal planes were analyzed to calculate the percentage of ERα+ and PR+ cells across the system. We found that ERα and PR responded differentially to EGF and FGF cues. EGF withdrawal increased the proportion of ERα+ cells, whereas FGF10 withdrawal reduced the percentage of ERα+ cells, and FGF2 withdrawal alone or in combination with EGF or FGF10 did not substantially alter ERα levels. In contrast, PR expression was selectively sensitive to FGF10. Withdrawal of EGF alone or FGF2 alone did not appreciably change the percentage of PR+ cells, but FGF10 withdrawal, either alone or combined with EGF, led to a marked reduction in PR+ cells, with a smaller decrease upon combined FGF2+EGF withdrawal. These patterns indicate that FGF10 plays a particularly prominent role in maintaining PR expression. These findings indicate that basal cell-derived organoids exhibit dynamic, heterogeneous hormone receptor expression that is responsive to both estrogen and microenvironmental cues, highlighting the plasticity of hormone receptor regulation in 3D culture. Thus, the integrated action of endocrine hormones and local growth factor signals dynamically shapes hormone-sensing states in mammary organoids, recapitulating key aspects of in vivo regulation of ERα and PR expression.

## Discussion

Hormone signaling in the mammary epithelium is typically interpreted through bulk measurements of receptor abundance and downstream transcriptional output. However, whether receptor expression reliably predicts hormone responsiveness at the level of individual epithelial cells has remained unclear. Using primary murine mammary organoids analyzed at single-cell resolution, we show that hormone receptor expression and hormone-driven transcriptional programs are intrinsically heterogeneous across individual cells. This heterogeneity reflects the combinatorial influence of receptor abundance, transcriptional co-regulator balance, and cellular state, rather than simple scaling with receptor levels. Importantly, this variability persists in organoid culture where microenvironmental complexity is reduced, indicating that heterogeneous hormone responses are not solely a consequence of tissue architecture or tumor-associated alterations. By quantifying the fraction of hormone-responsive genes activated in individual cells, we show that variability in transcriptional co-regulator abundance shows a stronger association with endocrine response magnitude than receptor transcript levels alone. This analysis reveals that variability in co-regulator abundance is not limited to permitting ER and PR transcriptional activity but is associated with the magnitude of hormone-responsive gene activation in individual epithelial cells.

Both basal cell-derived and ZsGreen⁺/PROM1⁺ organoids generate luminal hormone receptor-positive cells, yet they differ substantially in the relative abundance of ERα and PR. *Esr1* transcript levels are comparable across these systems. However, despite similar ERα expression, ZsGreen⁺/PROM1⁺ organoids display attenuated estrogen-induced transcription, suggesting that regulatory context rather than receptor abundance determines signaling strength. In contrast, Pgr expression is markedly higher in basal cell-derived organoids, consistent with prior reports demonstrating context-dependent regulation of PR in the mammary epithelium^63–65^. Because *Pgr* is an estrogen-responsive gene, differences in baseline *Pgr*^+^ fractions may reflect distinct prior signaling histories or regulatory states.

Progesterone responses further emphasize the role of co-regulator balance in shaping hormone signaling output. In basal cell-derived organoids, progesterone induces a bimodal transcriptional response, indicating that only a subset of PR-expressing cells mount a robust response. This behavior coincides with pronounced single-cell variability in the expression of transcriptional co-regulators, including both co-activators (e.g., *Src-1*) and co-repressors (e.g., *Ncor2*). Within this co-repressor family, *Ncor2* expression showed a significantly stronger association with single-cell progesterone response magnitude than *Ncor1*, suggesting that closely related co-repressors can contribute unequally to acute hormone signaling in mammary epithelial cells. This interpretation is consistent with prior work demonstrating non-redundant roles for NCOR1 and NCOR2 in modulating nuclear receptor-dependent transcriptional programs in breast and other hormone-responsive tissues^66,67^. In addition to co-regulator variability, receptor abundance itself is dynamically modulated by microenvironmental cues. Niche perturbation experiments reveal that growth factors differentially regulate ERα and PR protein levels, with FGF10 exerting effects distinct from EGF in basal cell-derived organoids. In the context of 3D mammary organoids, this suggests that local growth factor and paracrine signaling environments can tune ER/PR responsiveness across neighboring cells. Together, these findings indicate that endocrine phenotype emerges from the integration of intrinsic regulatory variability and extrinsic niche signals that shape receptor availability and signaling capacity.

The context-dependent relationship between ERα and PR expression observed in organoids closely parallels clinical observations in human breast cancer. PR status is routinely used to refine luminal subtype classification and to inform prognosis and endocrine therapy responsiveness^68–72^. ER⁺/PR⁻ tumors differ from ER⁺/PR⁺ tumors in clinicopathologic features and clinical outcomes and often derive less benefit from endocrine therapy despite ERα expression^69,71,73^. Mechanistic studies have implicated altered ER signaling and pathway rewiring in this reduced endocrine sensitivity^74^. Consistent with this clinical paradigm, our organoid data show that when PR expression is low, ERα remains detectable, yet estrogen-driven transcription is attenuated. These findings provide a mechanistic rationale for understanding how discordant ER/PR states emerge and how endocrine sensitivity may be reshaped independently of ERα abundance^75–78^.

At the single-cell level, heterogeneity likely reflects mixtures of cell states within each model system. In organoids, variability in baseline co-regulator expression, cell cycle status, chromatin accessibility, and lineage identity may all contribute to divergent hormone responses. Consistent with this interpretation, our ZsGreen⁺/PROM1⁺ sorted population was enriched for mature luminal signatures rather than progenitor states. This difference may reflect developmental stage differences in PROM1+ populations, as our study used adult mice whereas prior studies have primarily characterized ERα+/PROM1+ cells during puberty^29^. In contrast, heterogeneity in MCF7 cells may be further shaped by asynchronous cell cycling, genomic instability, and cell-intrinsic differences in chromatin accessibility that give rise to divergent transcriptional states, as reported in single-cell studies of ER⁺ breast cancer models^79–81^. Despite these differences in origin and regulatory architecture, both systems exhibit pronounced hormone response heterogeneity, suggesting that variable endocrine signaling is an intrinsic property of estrogen-responsive cells.

While heterogeneity is shared across systems, response kinetics differ markedly. scRNA sequencing reveals that basal cell-derived organoids mount rapid estrogen responses, with the fraction of estrogen-responsive genes per cell peaking within four hours and plateauing by sixteen hours. In contrast, MCF7 cells display a more gradual increase in estrogen-responsive gene fractions over later time points, consistent with previously described multi-phase estrogen responses in breast cancer cell lines^82–84^. These findings indicate that tissue context influences not only whether individual cells respond to hormone stimulation, but also the speed and coordination of those responses^56,85,86^. The faster and more synchronized responses observed in organoids likely reflect features of tissue context that are poorly preserved in 2D cell culture, including three-dimensional architecture, local microenvironmental signaling, and lineage-specific chromatin landscapes. Consistent with this interpretation, multiple studies have shown that breast organoids and other 3D culture systems more faithfully maintain primary tissue organization, receptor expression patterns, and treatment responses than monolayer cell lines^22,59,87–89^.

Hormone signaling in mammary epithelium therefore reflects the regulatory capacity of individual epithelial cells rather than receptor abundance alone. This framework helps explain why endocrine responses can vary across mammary epithelial populations despite similar receptor expression and provides a mechanistic basis for the heterogeneous transcriptional responses observed in both normal mammary tissue and hormone-responsive breast cancers. Understanding how endocrine signaling output emerges from the integration of intrinsic regulatory capacity and microenvironmental signaling therefore has important implications for mammary gland biology and for interpreting variability in clinical responses to endocrine therapies. By enabling controlled interrogation of hormone responses at single-cell resolution, mammary organoids provide a tractable platform to dissect how transcriptional co-regulators and niche-derived signals shape endocrine signaling programs in both normal mammary epithelium and malignant contexts. Future studies leveraging this system may help define how specific co-regulatory networks and microenvironmental signals interact to modulate endocrine responsiveness across epithelial cell states.

## Supporting information

Table S1 Cell Identity Statistics

Table S2 Bulk RNAseq Differential Expression

Table S3 SingleCell Hormone Response Statistics

Table S4 Hormones Supplemets

## Acknowledgements

We thank the members of the Rodriguez Laboratory and the NIEHS Epigenetics and RNA Biology Laboratory for ongoing support, insightful discussions, and constructive feedback. We are grateful to Brian Papas and the NIEHS Integrative Bioinformatics Support Group for bioinformatics support and guidance. We thank Jeff Tucker and Erica Scappini of the NIEHS Fluorescence Microscopy and Imaging Center for technical assistance with imaging. We also acknowledge Maria Sifre and Carl Bortner of the NIEHS Flow Cytometry Center for their support with cell sorting experiments. We thank Jason Malphurs and Xin Xu of the NIEHS Epigenomics Core Laboratory for next-generation sequencing expertise. Finally, we thank Mesut Muyan (Middle East Technical University) Paul Wade (NIEHS/NIH) and Carmen Williams (NIEHS/NIH) for critical review of the manuscript and data.

## Author Contributions

Conceptualization: P.Y. and J.R., Methodology: P.Y., Investigation: P.Y., Formal analysis: P.Y., C.R.D., B.D.B., J.A.H., and J.R., Resources: L.G.K., Data curation: P.Y., Writing-original draft: P.Y., Writing-review & editing: P.Y., C.R.D., J.A.H., T.K.A. and J.R., Supervision: J.R.

## Data Availability

The datasets generated in this study have been deposited in the Gene Expression Omnibus (GEO) under accession number GSE325083.

## Method details

### Experimental model and subject details

ERα-ZsGreen reporter mouse line was a kind gift from Yong Xu (Baylor College of Medicine, Children’s Nutrition Research Center, Texas, USA). C57BL/6J mice were obtained from The Jackson Laboratory. All animal experiments conform to Guide for the Care and Use of Laboratory Animals and were approved by the Institutional Animal Care and Use Committee at the National Institute of Environmental Health Sciences.

### Flow Cytometry

Abdominal and inguinal mammary glands were dissected from 8- to 12-week-old female mice. Dissociation mixture is prepared using the complete media composed of DMEM/F12 media (Gibco 11039-021), 5% Fetal Bovine Serum (CAT#), and 0.1% Gentamicin (Sigma G1397). 50 mg Collagenase (Type 4 filtered, Worthington LS004212) and 3 kunitz units Hyaluronidase (Purified, Worthington LS005475) are used to prepare 30 ml of dissociation mix. Dissected tissue is placed into pre-warmed (37°C) dissociation mixture and incubated on shaker at 400 rpm for 2 hours. During the incubation the suspension was pipetted every 30 min. After the incubation cells should be uniformly distributed into the mix and no visible tissue chunks were left. At the end of the incubation cells were centrifuged at 350 g for 5 min and the supernatant was discarded. Cells were resuspended in 5 ml of red blood cell lysis buffer and pipetted for at least 10 times. Hanks’ Balanced Salt Solution (HBSS) was supplemented with 2% FBS (HF) for the washes and buffer preparation. Red blood cell lysis buffer (10X, BioLegend 420301) was diluted in HF buffer. After the lysis cells were centrifuged at 350 g for 5 min and supernatant was discarded. Cells were resuspended in 5 ml of pre-warmed (37°C) TrypLE Express Enzyme (Gibco 12604021) and mixed by pipetting for at least 10 times. 10 ml of HF buffer is added, and cells were centrifuged at 350 g for 8 min and supernatant was discarded. Cells were resuspended in 2 ml of pre-warmed (37°C) HBSS and ∼ 500 kunitz units of DNase I (Worthington LK003172) was added and mixed by pipetting. 10 ml of HF buffer was added, and cells were filtered through 40 um filter (Falcon 352340). Cells were centrifuged at 350 g for 8 min and supernatant was discarded. If the pellet is heavily contaminated with red blood cells, lysis step was repeated. Flow staining buffer (PBS, 2.5 mM EDTA, 1% BSA, 3% FBS) and flow blocking buffer (PBS, 2.5 mM EDTA, 1% BSA, 10% FBS) were prepared. Cells were resuspended in 1 ml of flow blocking buffer and transferred into flow cytometry tubes (Falcon, 352058) and incubated on ice for 15 min. During this incubation, cells were counted, and antibody staining performed at 25 µl antibody mix per 10x106 cells for 40 min on a racker at 4°C. For ZsGreen+/Prom1+ cell sorting, a CD133 antibody was used. For basal and luminal cell sorting TER-119, CD31, and CD45 antibodies were used as dump channel, and cell surface markers CD24 and CD29 were used to distinguish luminal and basal populations. After staining, cells were centrifuged at 350 g for 5 min and resuspended in staining buffer at 10-20x106 cells/ml in the flow cytometry tube and sorting was carried out on BD Symphony S6 or BD FACSAriaII following gating and sorting strategies described in the previous study^30^. Sorted cells were collected into tubes pre-coated with 20% FBS to minimize cell loss and preserve viability.

### Organoid Culture

Single cells were seeded in basement membrane extract (BME; GelTrex) as small domes in uncoated 24-well plates (40 μl of cell-Basal Organoid Media (BOM) (60% GelTrex) per well, containing 104 cells) and cultured in DMEM/F12 supplemented with factors listed in Table S4 for 15-21 days. Organoids were passaged once using TrypLE Express to dissociate into single cells, which were counted, and 8x103 single cells were reseeded in fresh GelTrex. This first passage allowed expansion of organoids in culture to obtain sufficient material for downstream experiments. Hormone treatments were performed after maintaining organoids in same medium without refreshing for 3 days; at that point, complete medium (COM) was replaced with COM containing 17β-Estradiol (E2; 1μM for bulk RNA-seq, 10 nM for scRNA-seq and imaging), Progesterone (P4, 2.5 μM for bulk RNA-seq, 1 μM for scRNA-seq and imaging), or equivalent concentration of molecular-grade ethanol as vehicle control. For hormone depletion studies, at the time of medium change, organoids were refreshed with COM lacking EGF only, FGF2 only, FGF10 only, or specified combinations thereof.

### Single Cell RNA Sequencing

Organoid cells were dissociated using TrypLE Express to obtain single-cell suspensions for scRNA-seq and sorted mouse mammary cells were processed immediately after sorting since they are already at the single-cell suspension. Single cell suspensions from the hormone treated cells were also obtained using TrypLE Express after E2, P4 or vehicle control treatment for 4h or 16h. The cells were counted and examined for viability with AO/PI staining using a Luna FX7 cell counter (Logos Biosystems). About 8,500 live cells each sample at the concentration of 1.4-1.6 x10^6 cells/ml with viabilities more than 90% were loaded into the Single Cell Chip to generate single cell emulsion in Chromium X with Chromium GEM-X Single Cell 3’ Kit v4 (10x Genomics, Cat. 1000686). Reverse transcription of mRNA and cDNA amplification were carried out following the manufacture’s instruction (10x Genomics, CG000731). The amplified cDNA was further fragmented to construct NGS libraries. The libraries were then sequenced by the NIEHS Genomics Core Laboratory with the parameters recommended in the manufacture’s instruction. > 200 million raw reads were obtained for each library. Sequencing data generated by the 10X libraries were processed into count matrices using cell ranger v9.0.1, with gene quantification performed against the gex-GRCm39-2024-A reference transcriptome. Cells with mitochondrial gene percentage > 10% were removed. Downstream single-cell analyses were performed in R using the Seurat package (v5.0.1), following the standard workflow for normalization, dimensionality reduction, and graph-based clustering. Cell cycle scores were determined for all cells using the CellCycleScoring function in the Seurat package (v5.0.1), after mapping human cell cycle gene sets from cc.genes.updated.2019 to their mouse orthologs using the g:Profiler tool (via gorth() function in the gprofiler2 package). Lineage-associated transcriptional programs were quantified at single-cell resolution using module score–based lineage scoring. Gene sets representing basal, luminal progenitor, and mature luminal identities were defined from curated differentially expressed genes^32,33^ and scored for each cell using Seurat’s AddModuleScore function, which implements a Tirosh-style gene signature scoring strategy^90^. Module scores were independently rescaled to a 0-1 range to enable comparison across lineage programs. Normalized scores were visualized using ternary plots generated with the Ternary R package^91^, with each cell positioned according to its relative basal, luminal progenitor, and mature luminal scores and colored by Seurat cluster identity. To control differences in sequencing depth between datasets, UMI counts were normalized by downsampling. Raw RNA count matrices were extracted from Seurat objects, and the dataset with higher average UMIs per cell was downsampled to match the mean UMI depth of the lower-depth dataset using the downsampleMatrix function from the scuttle package^92^. Gene-positive cells were defined as those with at least one detected UMI, and percentages of Esr1-, Pgr-, Krt8-, and Krt14-expressing cells were calculated within organoid populations. Hormone-induced transcriptional responses in ERα+ and ERα- cells were quantified at single-cell resolution using a ratio of responding genes (RRG) framework adapted from previous study^55^. ERα+ cluster of cells were defined and rest of the cells identified as ERα- cells. DEGs were identified by comparing hormone-treated (E2 or P4) to matched vehicle (EtOH) samples at each time point using Seurat’s Wilcoxon based FindMarkers function, and for the RRG analysis we restricted to upregulated DEGs (positive log2 fold-change and passing the adjusted p-value threshold). For each upregulated DEG set, baseline log-normalized expression distributions were estimated from timepoint-matched vehicle ERα+ cells, computing per-gene means and standard deviations while excluding zeros. A gene was classified as “responding” in an individual hormone-treated cell (ERα+ and ERα-) if its expression exceeded the baseline mean plus one standard deviation, and the RRG score for each cell was defined as the proportion of DEGs meeting this criterion. RRG distributions for hormone-treated and vehicle groups were then compared using Wilcoxon rank-sum tests and visualized as percentages of responding genes per cell. All single-cell visualizations (including UMAPs, dot plots, violin plots, boxplots, and volcano plots) were generated in R using Seurat and the ggplot2 package (v4.0.1).

### Cleavage under targets and tagmentation (CUT&Tag)

CUT&Tag was performed following the Bench top CUT&Tag V.3 protocol^54^. Briefly, we used 50K nuclei isolated from basal organoid and basal sorted and luminal sorted cells. 10 μL BioMag-Plus Concanavalin (Bangs Laboratories) were used per reaction to immobilize the nuclei. Then we added 1:50 rabbit H3K27ac (Active Motif), 1:50 rabbit H3K27me3 (Cell Signaling Technology antibody and incubated for 2h. After one hour of secondary antibody incubation 1.25μL of CUTANA pAG-Tn5 adapter complex was used to load the enzyme to the antibody bound regions. One hour of tagmentation at 37°C was followed by DNA extraction using MinElute PCR Purification Kit (Qiagen). Extracted DNA was subjected to PCR amplification using unique primers sets (Nextera XT v2 Full set (N7-S5)). We amplified both antibody samples 15 PCR cycles. Libraries were size selected using AMPure XP Beads (Beckman Coulter) with 1.3X ratio. We quantified and assessed the quality of the libraries with Qubit Flex and Tapestation and then sequenced the libraries with 50 bp paired-end reads on Illumina high output NovaSeq SP.

CUT&Tag reads were processed using pipelined adapted from^78,93^. First adaptors were trimmed using Cutadapt and reads were aligned to the mm10 genome assembly using Bowtie2. Samtools was used to sort the reads and deduplication was performed using Picard Tools (http://broadinstitute.github.io/picard). Coverage was generated using bamCoverage and RPKM normalization. Peaks were called using MACS2 and filtered to keep peaks that were called in more than one replicate of at least one condition. Peaks were quantified with the featureCounts tool and differential analyses was performed using the DESeq2 R package.

### Bulk RNA sequencing

Total RNA was isolated from freshly sorted ZsGreen+/Prom1+ and basal cells. Total RNA was also isolated from the ZsGreen+/Prom1+ or basal organoids that are treated with E2. P4, E2+P4 or vehicle control for 4h and 24h. Total RNA was isolated using TRIzol Reagent by adding directly the solution onto the wells. General TRIzol protocol was followed. Briefly, 400 µl of TRIzol Reagent was directly added onto the organoid domes in the 24-well plate and by pipetting the basement membrane is become soluble and the content was transferred into a Nuclease-free Eppendorf tube. Samples were incubated at room temperature for 3 minutes to allow complete dissociation of the nucleoproteins complex. 120 µl of chloroform was added to each sample and tubes were upside down for 40 times to mix. After that incubated at room temperature for 3 minutes. Then samples were centrifuged at 11000 x g at 4°C for 15 min. The upper aqueous phase containing the RNA is transferred into a new Nuclease-free Eppendorf tube. 1 µl Glycogen RNA-grade (Thermo Scientific) and 200 µl isopropanol added and incubated at -20°C overnight for RNA precipitation. Next day, samples were centrifuged at 11000 x g at 4°C for an hour. Pellet was washed with 100 µl of 75% ethanol and centrifuged at 11000 x g at 4°C for 15 min. Wash step repeated two times. RNA pellet was then allowed to dry at room temperature for 10 minutes and resuspended in 20 µl of Nuclease-free water. The quantity and quality of the RNA samples were assessed by Tapestation. Polyadenylated mRNAs were selected using Dynabeads Oligo(dT)25 (ThermoFisher Scientific) and libraries were prepared with NEBNext Ultra Directional RNA Library Prep Kit for Illumina (NEB) following manufacturer’s instruction. Libraries were size selected using SPRIselect beads (Beckman Coulter). Libraries were quantified Qubit Flex and Tapestation and then sequenced with 75 bp single-end reads on Illumina NextSeq high-output. Reads were filtered so that only those with a mean quality score of 20 or greater were kept. Adapter was trimmed using Cutadapt version 3.7. Reads were aligned to the hg38 genome assembly using STAR version 2.6.0c. Counts were obtained using the featureCounts tool from the Subread package version 1.5.1 with the GENCODE basic gene annotation version 44. Differential expression was quantified using the DESeq2 R package version 1.34.0.

### Immunocytochemistry (ICC)

Whole-mount immunofluorescence staining was performed on intact organoids recovered from basement membrane extract. Organoids were first released from GelTrex domes using Gentle Cell Dissociation Reagent (STEMCELL Technologies) according to the manufacturer’s recommendations for breaking up Matrigel-like matrices. Briefly, culture medium was aspirated and 500 µl Gentle Dissociation Reagent was added per well of a 24-well plate. Domes were disrupted by triturating twice with a 1 ml pipette tip that had been cut to widen the bore, and the resulting suspension was transferred to 1.5 ml microcentrifuge tubes. Tubes were placed on ice on a rocking platform and incubated for 20 min to dissolve the matrix. Organoids were then pelleted by centrifugation at 600 g for 6 min. For fixation, the supernatant was aspirated and organoids were resuspended in 300 µl of 4% paraformaldehyde (PFA) in PBS and incubated for 45 min at room temperature on a rocking platform on a rack. Organoids were pelleted by centrifugation at 600 g for 6 min, washed once with 300 µl Immunofluorescence (IF) buffer (PBS supplemented with 2% BSA, 0.2% TritonX-100, and 0.05% TWEEN20), and centrifuged again at 600 g for 6 min. When necessary, fixed organoids were resuspended in PBS and stored at 4-8°C, although staining was typically continued immediately to minimize loss of antigenicity. Antigen retrieval was performed by resuspending organoids in 200 µl pre-warmed citrate buffer (10 mM Sodium Citrate buffer with 0.05% Tween 20 (pH 6.0)) and incubating the tubes in a 98°C heating block for 20 min. The heating block was then switched off and organoids were allowed to cool in place for an additional 20 min. Samples were pelleted by centrifugation for at 600 g 6 min and the supernatant was removed. To quench residual aldehydes, organoids were resuspended in 300 µl of 0.3 M glycine in PBS and incubated with gentle rocking for 30 min at room temperature, followed by centrifugation at 600 g for 6 min. For permeabilization and blocking, organoids were resuspended in 300 µl permeabilization/blocking solution consisting of permeabilization buffer (1% Triton X-100 in PBS [v/v]) supplemented with 5% normal goat serum (v/v), chosen to match the host species of the secondary antibodies. Organoids were incubated in blocking solution overnight at 4°C. After blocking, organoids were washed three times in IF buffer, with centrifugation at 600 g for 6 min between washes. Primary antibody staining was performed by resuspending organoids in 200 µl of IF buffer containing mouse anti ERα (1:100, Abcam), rabbit anti-PR (1:100), and rat anti-keratin 8 (K8; 1:100; TROMA-I was deposited to the DSHB by Brulet, P. / Kemler, R. (DSHB Hybridoma Product TROMA-I)). Samples were incubated overnight at 4°C with gentle rocking (extended incubations over a weekend did not noticeably alter staining intensity). Following primary incubation, organoids were centrifuged at 600 g for 6 min, the primary antibody solution was aspirated, and organoids were washed three times in IF buffer as above. For secondary antibody staining, organoids were resuspended in 200 µl of IF buffer supplemented with 10% normal goat serum and incubated with goat anti-mouse Alexa Fluor 594 (Thermo Fisher), goat anti-rabbit Alexa Fluor 488 (Thermo Fisher), and goat anti-rat Alexa Fluor 750 (Abcam) secondary antibodies (1:500). Samples were incubated overnight at 4°C with gentle rocking (3-4 h incubations at room temperature were also sufficient and gave comparable staining). Organoids were then washed three times in IF buffer, pelleting at 600 g for 6 min between washes. Nuclei were counterstained by adding DAPI directly to the final secondary antibody or wash solution to a final concentration of 2-4 µg/ml and incubating organoids for 20 min at room temperature in the dark. Organoids were centrifuged at 600 g for 6 min, washed once with water and once with PBS, with 6 min centrifugation at 600 g after each wash. For mounting, organoids were resuspended in 40-50 µl Fluoromount-G and transferred onto glass coverslips for imaging.

### Imaging

Confocal images of single-organoids were taken on a Zeiss LSM980 with Airyscan2 (Carl Zeiss Inc, Oberkochen, Germany) using a Plan-Apochromat 40X/1.3 Oil DIC objective. A 730nm DPSS laser at 1% power was used to excite Alexa750 while emissions were collected using a 755-900nm bandpass filter. Alexa594 was excited using a 561nm DPSS laser at 3% power and emissions were filtered with a 578-649nm bandpass filter. A 488 nm laser at 3% power was used for excitation of Alexa488 while a 498-569nm bandpass filter was used to collect the emission signal. DAPI was excited with a 405nm Diode laser at 0.5% and a 408-474nm bandpass filter collected the emission. Each channel was acquired sequentially, and the master gain of the detector optimized per channel (Alexa750: 500 V; Alexa594: 725 V; Alexa488: 725 V; DAPI: 500 V). Finally, all images were taken with a zoom of 1, a 1.02µs pixel dwell time, and a pixel size of 0.207µm. Z-stacks of the entire organoid were acquired with an Z-interval of 1µm.

For large field-of-view imaging, confocal tile-scan images were acquired on a Zeiss LSM980 with Airyscan2 (Carl Zeiss Inc., Oberkochen, Germany) using a Plan-Apochromat 10X/0.3 objective. Dyes were excited using the same laser lines described above (405 nm, 488 nm, 561 nm, and 730 nm) with laser powers set to 0.5% (405 nm), 3% (488 nm), 3% (561 nm), and 0.5% (730 nm). Emission signals were collected using the same bandpass filters described above with the standard confocal detector. Tiles were acquired with a 10% overlap to ensure accurate stitching, and the number of tiles (typically 30–50 per sample) was adjusted to cover the entire organoid area. Pixel dwell time was set to 1.02 µs, with a pixel size of 0.829 µm and a zoom factor of 1. Z-stacks were acquired with a Z-interval of 3 µm. Tile stitching was performed in ZEN 3.9 (Carl Zeiss Inc., Oberkochen, Germany) using default parameters and the DAPI channel as the reference channel.

## Image Analysis

### Individual organoid analysis

For single-organoid datasets, full 3D image stacks were analyzed. Nuclear segmentation was performed using Cellpose v3.0.11 with the DAPI channel as input, using the pretrained nuclei model with deblurring enabled. Segmentation was performed in 3D mode. Flow and cell probability thresholds were optimized to refine nuclei segmentation. Incomplete nuclei masks were filtered out with a size exclusion threshold. Using Python the median fluorescence intensities for ER and PR intensities were extracted across the 3D volume of nuclei masks. Nuclei positive for ER and PR were determined using thresholds, determined from on intensity histograms and refined by visual inspection of mask overlays to ensure accurate classification. Organoid-level values were calculated as the median per-cell intensity (or as fraction of positive nuclei within the organoid) within each organoid.

### Quantitative analysis of tile-scan images

For tile-scan datasets, three representative z-slices were extracted from the midpoint of the sample using FIJI (ImageJ). Nuclear segmentation was performed using Cellpose v3.0.11 with the DAPI channel as the input and the pretrained nuclei model with deblurring enabled. Flow and cell probability thresholds were used to refine segmentation and incomplete nuclei were removed using a minim size threshold. In Python nuclei masks were used to extract median fluorescence intensities for all channels. Histograms of single-cell intensities were generated to define initial thresholds to define ER and PR positive cells. The first intensity peak corresponded to background-level signal. Thresholds were iteratively refined by visual inspection of mask overlays. Summary statistics were calculated after marker classification, including (% positive cells, median intensity, organoid-level aggregation metric).

