## Supplementary Figures for "Layered Single-Cell Heterogeneity in Hormone Receptor Signaling Across Mouse Organoids and Human ERα+ Cancer Cells"

**SUPP. FIGURE 1**

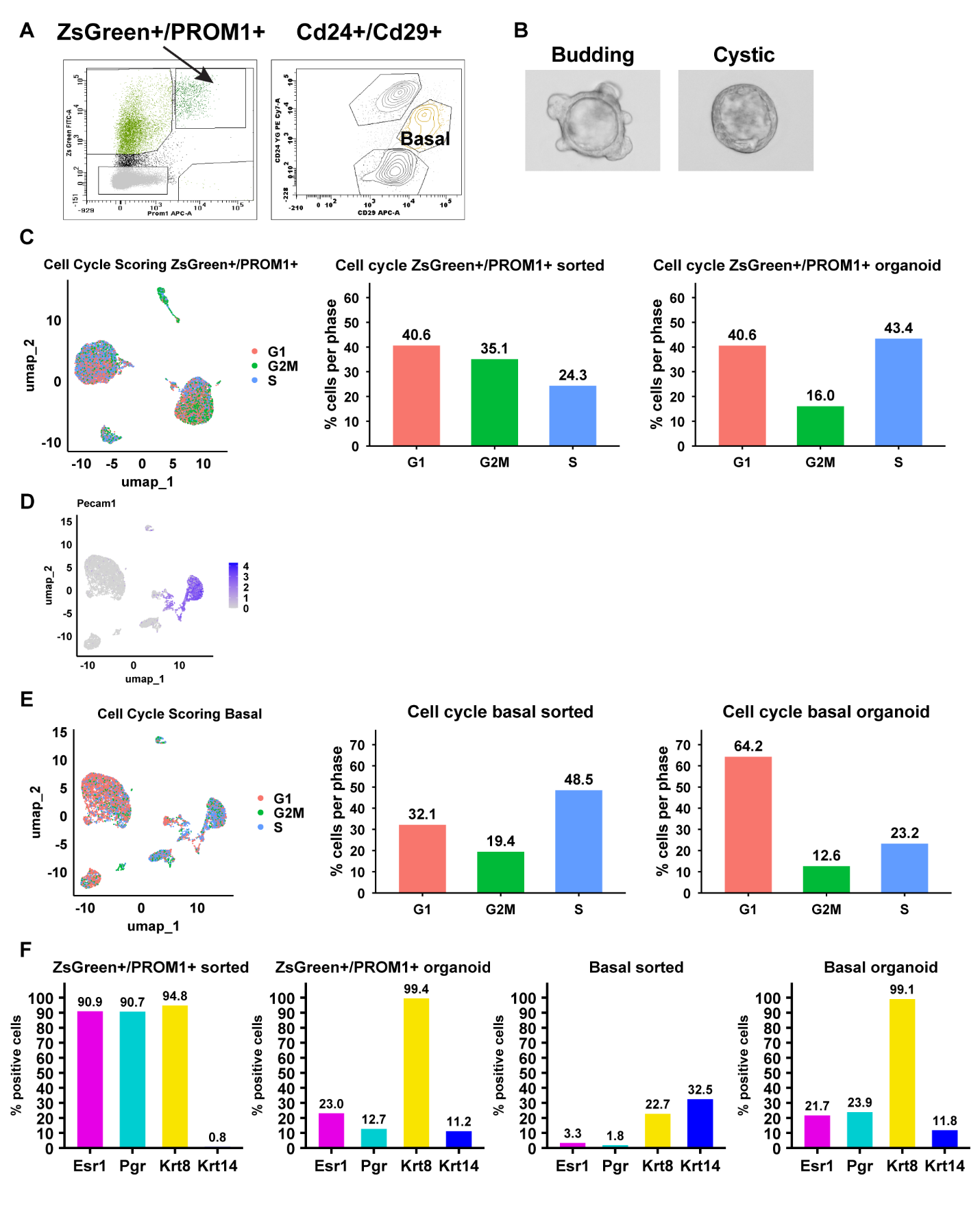

**Figure S1 Characterization of sorted populations, organoid morphology, and cell cycle distribution**

(A) Flow cytometry gating strategy for isolation of ZsGreen⁺/PROM1⁺ luminal-enriched cells and basal-enriched (CD24^+^CD29^hi^) epithelial populations from mouse mammary glands.
(B) Representative brightfield images of budding and cystic organoid morphologies observed in 3D culture.
(C) Cell cycle scoring of ZsGreen⁺/PROM1⁺-sorted cells and derived organoids. Left, UMAP colored by cell cycle phase; right, quantification of G1, S, and G2/M phase distribution.
(D) UMAP feature plot showing *Pecam1* expression, identifying a small endothelial-contaminated cluster retained during clustering analysis.
(E) Cell cycle scoring of basal-sorted cells and basal cell-derived organoids. Left, UMAP colored by cell cycle phase; right, quantification of cell cycle phase distribution.
(F) Percentage of Esr1^+^, Pgr^+^, Krt8^+^, and Krt14^+^ cells in sorted and organoid populations derived from ZsGreen⁺/PROM1⁺ and basal cells.

**SUPP. FIGURE 2**

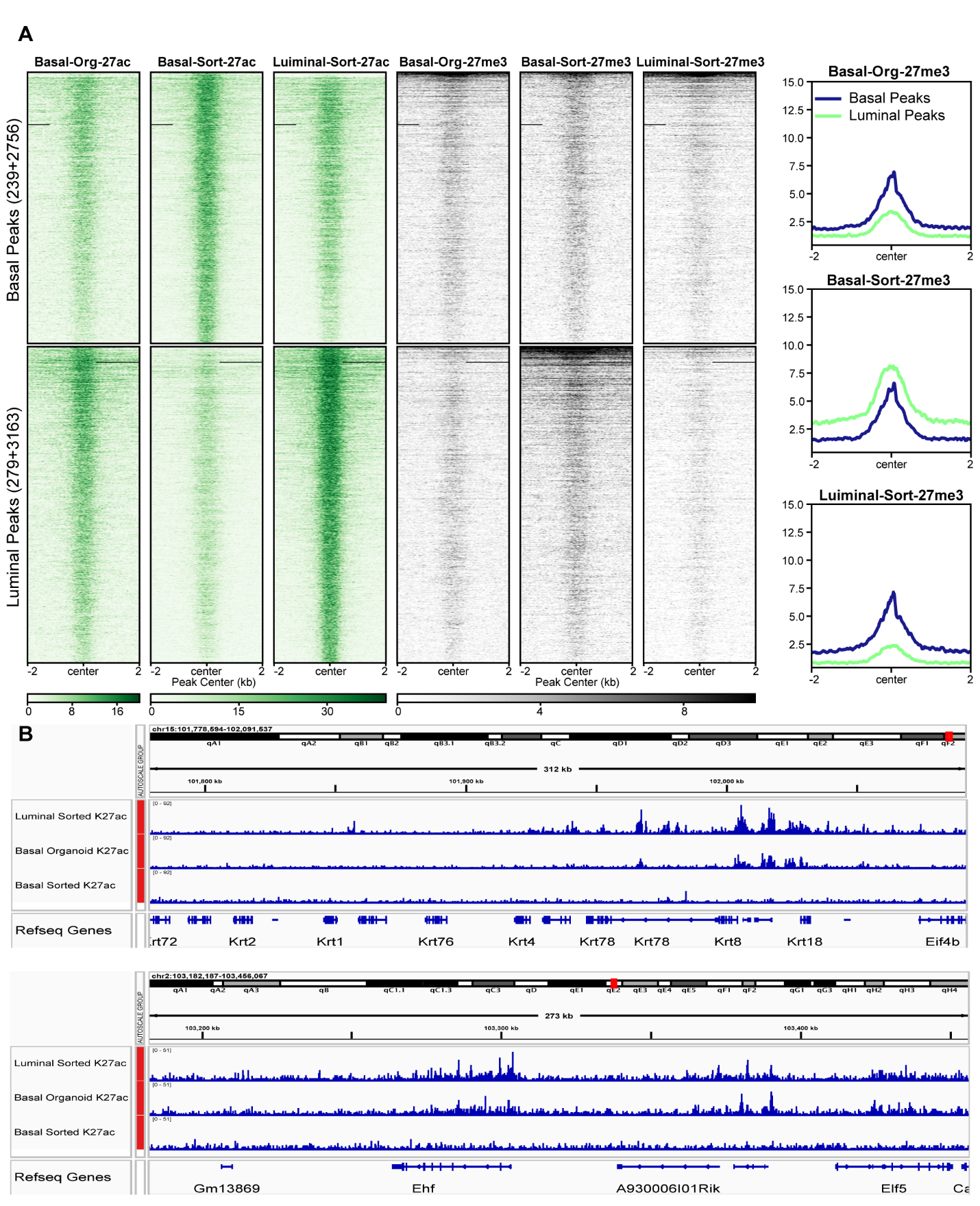

**Figure S2 H3K27ac and H3K27me3 profiles at lineage-associated peaks**

(A) Heatmaps of H3K27ac and H3K27me3 CUT&Tag signal centered on basal-associated peaks (top) and luminal-associated peaks (bottom) across basal cell-derived organoids, basal-sorted, and luminal-sorted epithelial populations. Signal is shown ± 2kb from peak centers. Right panels show aggregate signal profiles of H3K27me3.
(B) Genome browser tracks showing H3K27ac CUT&Tag signal at the Krt8 and Elf5 loci across luminal-sorted, basal-sorted cells and basal cell-derived organoids. Signal is displayed as normalized coverage tracks aligned to mm10 genome.

**SUPP. FIGURE 3**

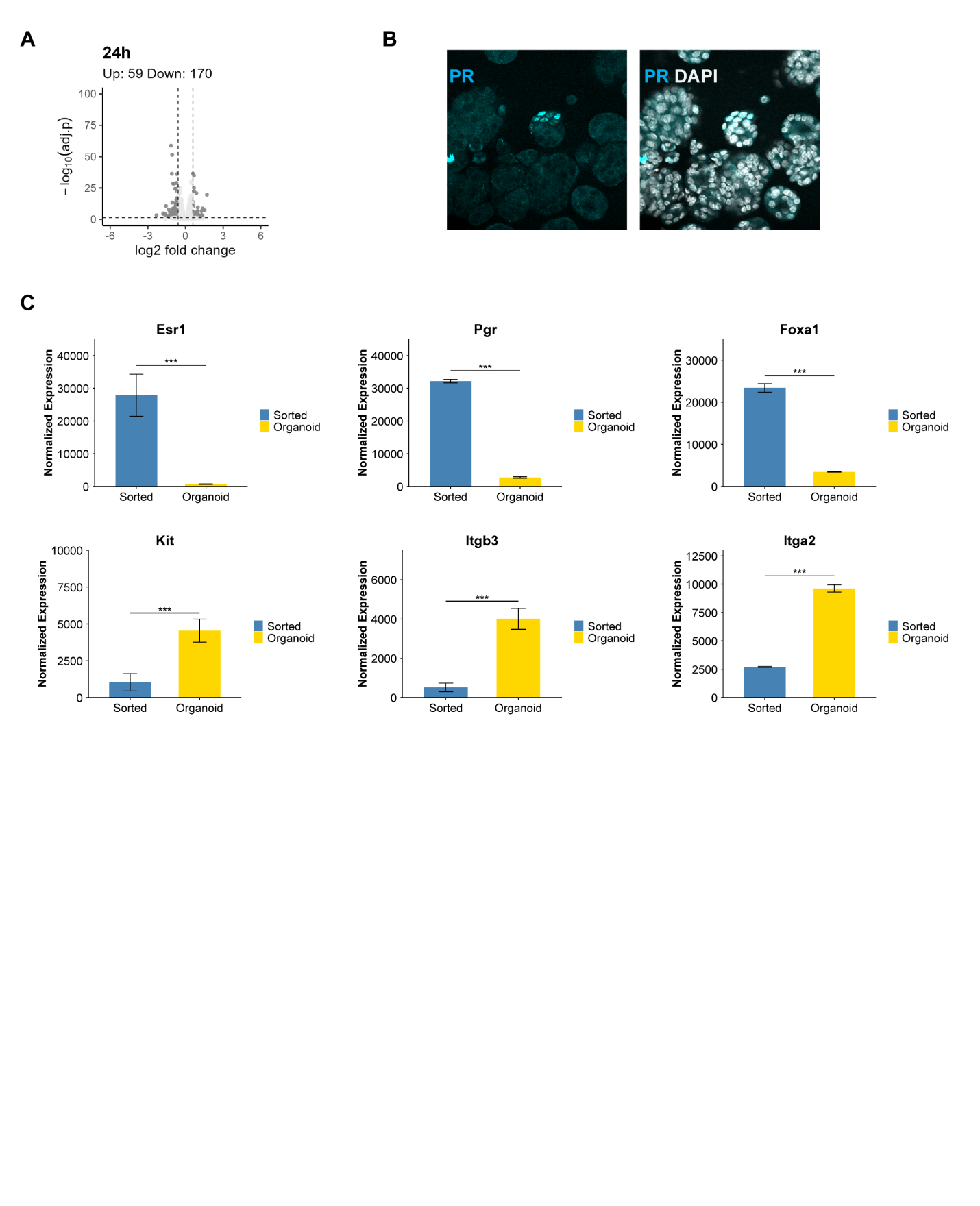

**Figure S3 Reduced receptor expression and hormone response in ZsGreen⁺/PROM1⁺ organoids**

(A) Volcano plot of differentially expressed genes in ZsGreen⁺/PROM1⁺ organoids at 24h following E2 treatment relative to vehicle control. Differential expressionwas defined as adjusted p value < 0.05 and |log2 fold change| ≥ 0.585.
(B) Representative confocal images of PR immunostaining (cyan) in ZsGreen⁺/PROM1⁺ organoids. DAPI marks nuclei (gray).
(C) Bulk RNA-seq expression of Esr1, Pgr, Foxa1, Kit, Itgb3, and Itga2 in freshly sorted ZsGreen⁺/PROM1⁺ cells and derived organoids. Bars represent mean ± SD of normalized counts. Statistical significance reflects DESeq2 differential expression analysis.

**SUPP. FIGURE 4**

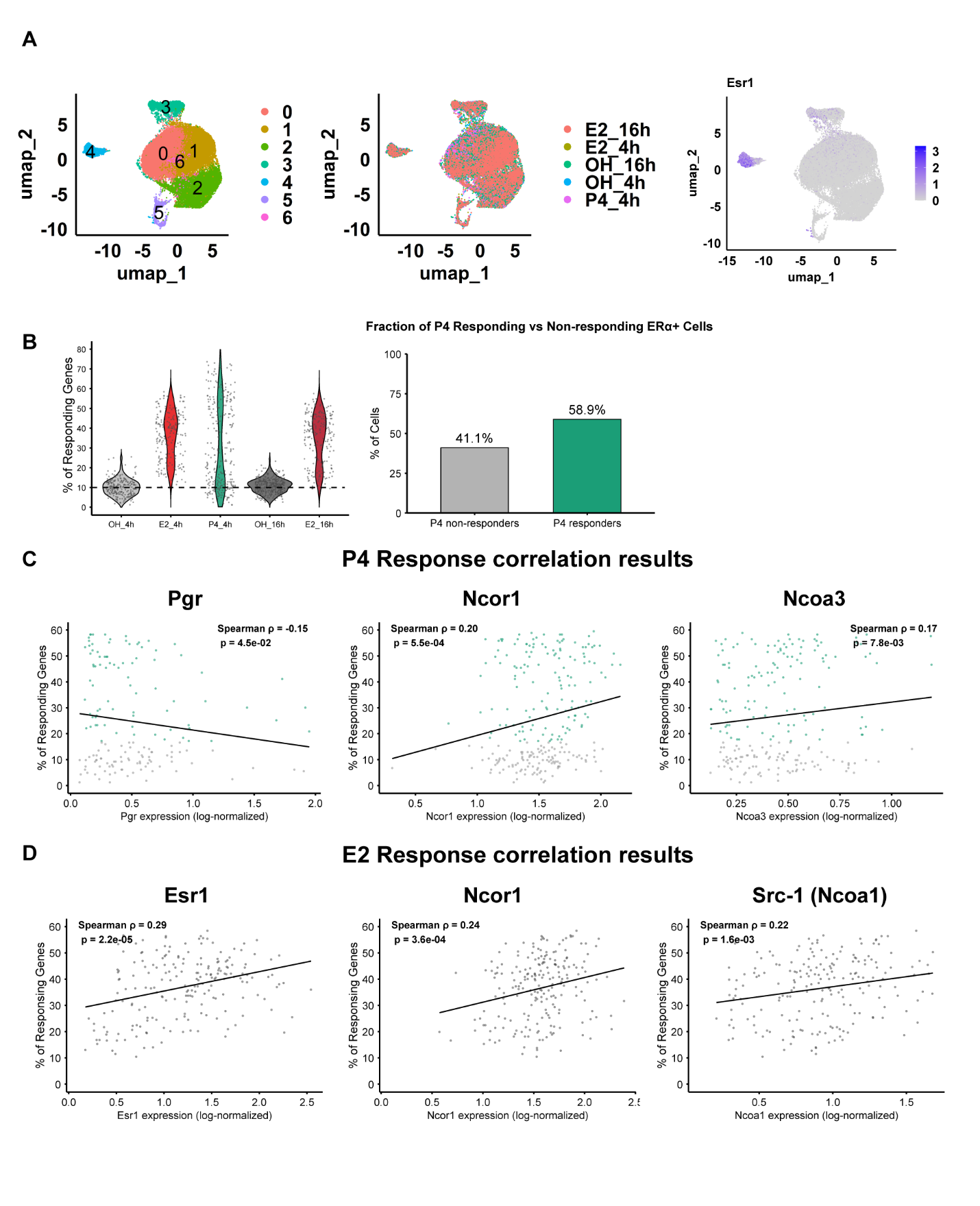

**Figure S4 Single-cell analysis of progesterone response heterogeneity in basal cell-derived organoids**

(A) UMAP representation of basal cell-derived organoid cells following vehicle (EtOH), E2 (10nM), or P4 (1 µM) treatment for 4h and 16h. Left panel shows unsupervised clustering; right panel shows distribution by treatment and time point.
(B) Violin plots showing the percentage of responding genes (%RRG) per ERα^+^ cells across conditions. The dashed line indicates the median baseline %RRG in ERα^-^ cells. Right panel quantifies the proportion of ERα^+^ cells classified as P4 responders versus non-responders.
(C) Scatter plots showing the relationship between Pgr transcript abundance (log-normalized) and P4-induced %RRG in ERα^+^ cells. Spearman correlation coefficient (ρ) and p-value are indicated.

**SUPP. FIGURE 5**

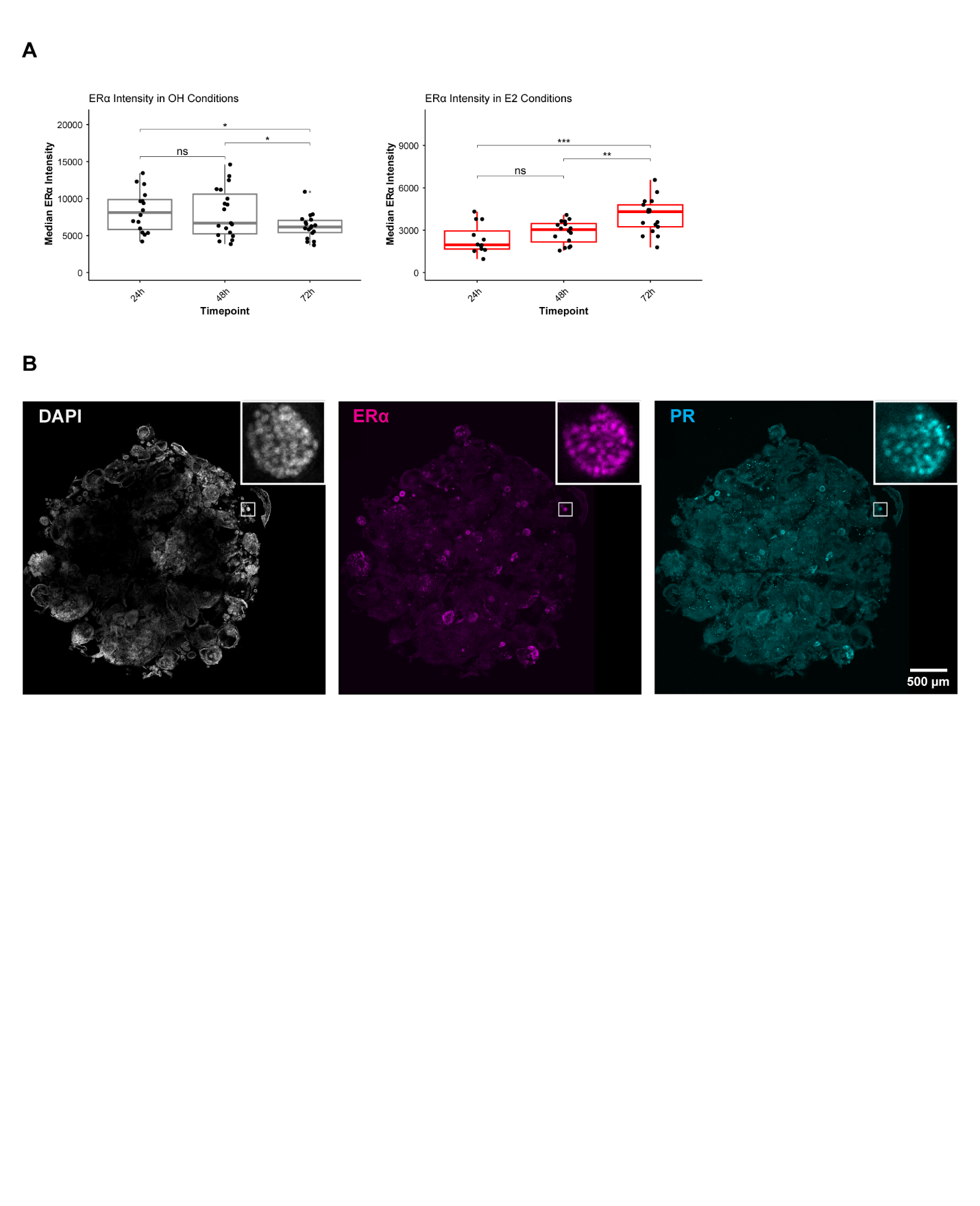

**Figure S5 Whole-well quantification and temporal analysis of ERα protein expression in basal cell-derived organoids**

(A) Quantification of ERα median fluorescence intensity in basal cell-derived organoids treated with vehicle (EtOH) or 10 nM E2 for 24h, 48h, 72h. In vehicle-treated organoids, ERα levels progressively declined over time, whereas E2-treated organoids exhibited increased ERα intensity at later time points, resulting in partial convergence or ERα levels by 72h. Statistical significance was determined using two-tailed unpaired Welch’s t-tests; pairwise comparisons are indicated.
(B) Representative confocal tile-scan images acquired using a 10x objective showing the entire well of basal cell-derived organoids stained for DAPI (gray), ERα (magenta), and PR (cyan). Insets display a single organoid extracted from the whole-well image to illustrate organoid morphology and receptor distribution. Tile-scan acquisition was used to capture all organoids within the well to minimize sampling bias for quantitative analysis. Scale bar, 500 µm.
